# Higher order thalamus encodes correct goal-directed action

**DOI:** 10.1101/2020.07.05.188821

**Authors:** D. LaTerra, S. Petryszyn, Marius Rosier, L.M. Palmer

## Abstract

The thalamus is the gateway to the cortex. Cortical encoding of sensory information can therefore only be understood by considering the influence of thalamic processing on sensory input. Despite modulating sensory processing, little is known about the role of the thalamus during sensory-based behavior, let alone goal-directed behavior. Here, we use two-photon Ca^2+^ imaging, patch-clamp electrophysiology and optogenetics to investigate the role of axonal projections from the posteromedial nucleus of the thalamus (POm) to the forepaw area of the primary somatosensory cortex (forepaw S1) during sensory processing and goal-directed behavior. We demonstrate that POm axons are active during tactile stimulus and increase activity specifically during the response and, to a lesser extent, reward epochs of a tactile goal-directed task. Furthermore, POm axons in forepaw S1 preferentially signaled correct behavior, with greatest activity during HIT responses. This activity is important for behavioral performance, as photoinhibition of archaerhodopsin-expressing neurons in the POm decreased overall behavioral success. Direct juxtacelluar recordings in the awake state illustrates POm neurons fire sustained action potentials during tactile stimulus. This tactile-evoked POm firing pattern was used during ChR2 photoactivation of POm axons in forepaw S1, revealing that action potentials in layer 2/3 (L2/3) pyramidal neurons are inhibited during sustained POm input. Taken together, POm axonal projections in forepaw S1 encode correct goal-directed active behavior, leading to GABA_A_-mediated inhibition of L2/3 pyramidal neurons.

## INTRODUCTION

Goal-directed behavior is crucial for survival in a dynamic environment. It involves the encoding and integration of sensory information that leads to specific rewarded behaviors (Kepecs et al., 2008; Li et al., 2015; Takahashi et al., 2016; Xu et al., 2012). The thalamus is a fundamental hub for the transfer of sensory information to the cortex, sending and receiving widespread innervation from numerous cortical and subcortical structures (Oh et al., 2014; Sherman and Guillery, 1996). Despite being perfectly positioned to coordinate the relay and integration of sensory information required during sensory-based behavior, historically, the thalamus has been viewed as a passive sensory relay center with negligible contribution to higher-order brain function and behavior. Recent studies have challenged this classical view, illustrating the thalamus plays crucial roles in cognitive tasks such as attention (Schmitt et al., 2017; Wimmer et al., 2015; Zhou et al., 2016), sensory perception (Saalmann and Kastner, 2011; Wilke et al., 2009), motor preparation and suppression (Casagrande et al., 2005; Yu et al., 2016), cortical plasticity (Gambino et al., 2014) and learning (Williams and Holtmaat, 2019).

The higher order thalamus is an enigmatic class of non-specific thalamic nuclei which send feedback input to sensory cortical areas (Sherman and Guillery, 1996). Specifically, the postero-medial nucleus of the thalamus (POm) is the higher order thalamic nucleus subtending sensory processing in the primary somatosensory cortex (S1) (Deschenes et al., 1998; Jones, 2007). Sending dense projections to layer 1 and layer 5 of S1(Meyer et al., 2010), the POm specifically targets a complex cortical microcircuit (Audette et al., 2017) which influences the encoding of somatosensory inputs (Mease et al., 2016; Urbain et al., 2015; Zhang and Bruno, 2019). POm is reciprocally connected with S1, but also receives and sends projections to motor, premotor, association cortices as well as many subcortical regions including the zona incerta and striatum (Alloway et al., 2017; Oh et al., 2014; Trageser and Keller, 2004; Yamawaki and Shepherd, 2015). Based on its influence on cortical sensory processing and known extensive connectivity, the POm may play an important role in behaviors which require both the perception and integration of sensory information, such as sensory-based goal-directed behavior. Since this behavior requires the integration of information from multiple brain regions, POm input onto layer 2/3 pyramidal neurons may play an important role as these neurons send and receive long-range projections from other cortical areas (Oh et al., 2014; Yamashita et al., 2018), making them the perfect target for the modulation of cortical information. To test this, we used whole-cell patch clamp and juxtacelluar electrophysiology, two-photon Ca^2+^ imaging and optogenetics to investigate the role of POm projections in the forepaw S1 during tactile goal-directed behavior.

## RESULTS

### Ca^2+^ imaging of POm axonal projections in forepaw S1 during tactile stimulation

Ca^2+^ imaging of POm axonal projections in forepaw S1 was performed in awake mice (P50 - 70) previously injected with the Ca^2+^ indicator GCaMP6f (AAV1.Syn.GCaMP6f.WPRE.SV40) into the POm (see Methods; Fig. 1A, and Figure S1). Following expression, Ca^2+^ transients were recorded from POm axons that project to layer 1 of the forepaw S1 (48 ± 6.8 μm from the pia surface; Figure S2). These POm axons were sparsely spontaneously active in awake naive mice, generating large Ca^2+^ transients (>2 s.d of the baseline fluorescence; see Methods) at 0.08 ± 0.01 Hz (n = 113 axons, 5 mice; Figure 1B). To test whether POm encodes tactile information, axonal Ca^2+^ transients were investigated during tactile stimulus of the contralateral forepaw (FP; 200 Hz, 500 ms). Here, POm axons significantly increased Ca^2+^ signaling above the spontaneous rate, evoking a response to 21.26 ± 2.22 % of forepaw stimuli (p < 0.0001; n = 113 axons, 5 mice; Figure 1C). Direct comparison of Ca^2+^ transients from POm axons with both spontaneous and evoked activity (n = 44 axons, 5 mice) shows there was no significant difference in the peak amplitude (1.15 ± 0.08 vs 1.12 ± 0.10 ΔF/F; p = 0.4078; Fig. 1D) nor duration (0.636 ± 0.878 vs 0.686 ± 0.142 s; p = 0.6761) of the Ca^2+^ transients evoked spontaneously or in response to tactile stimulus.

**Figure 1.**
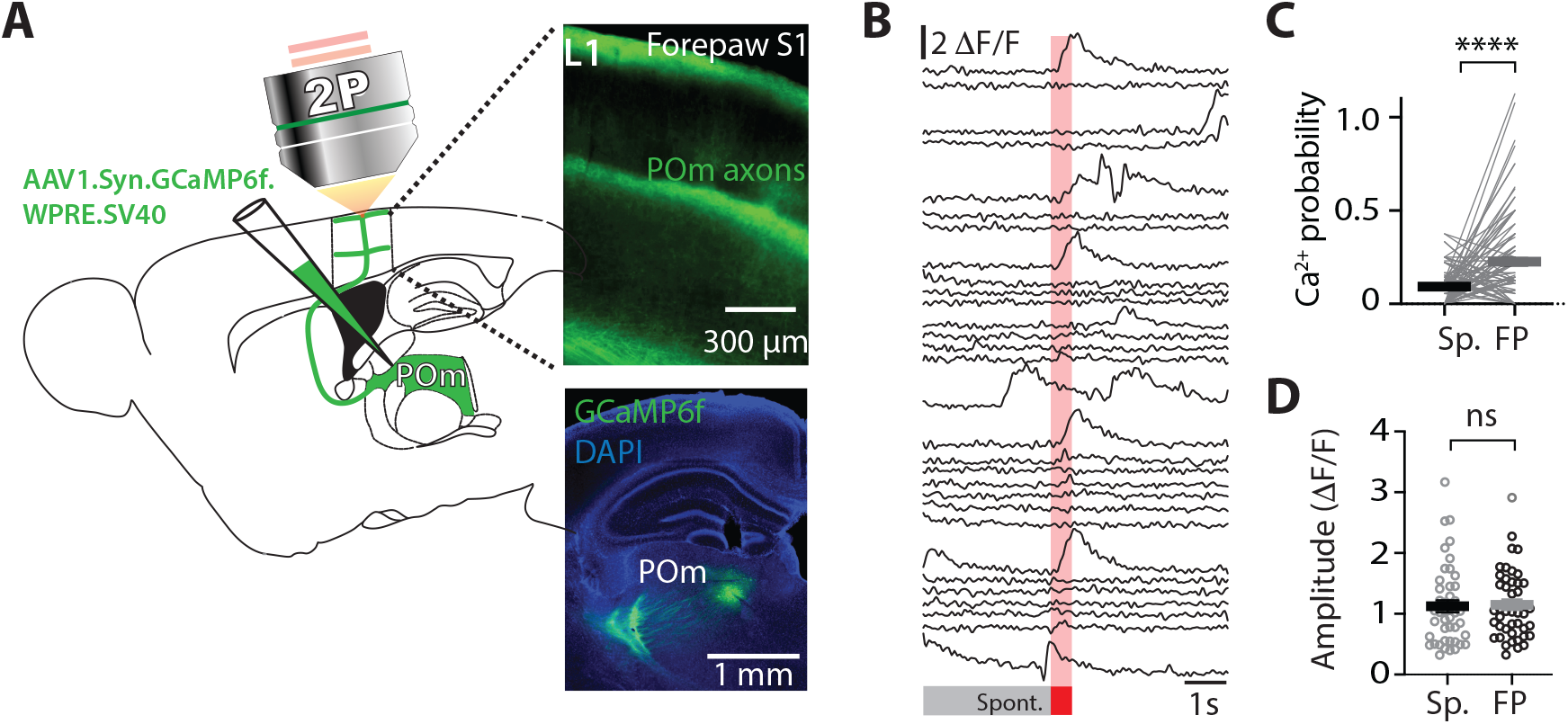
POm axonal projections in forepaw S1 encode sensory information. **(A**) Experimental design. The Ca^2+^ indicator GCaMP6f was locally injected into the POm and two-photon Ca^2+^ imaging was performed from POm axons projecting to layer 1 of forepaw S1 during tactile stimulation. Post hoc fluorescence images illustrating (top) that POm primarily sends axonal projections to layer 1 and layer 5 of the forepaw S1 and (bottom) localized POm injection. **(B)** Ca^2+^ traces from a representative POm axon in layer 1 of forepaw S1 in the naïve awake state during forepaw stimulation (FP; 200Hz, 500ms). **(C)** The probability of evoking a Ca^2+^ transient during forepaw stimulation (FP; black) versus spontaneous activity (SP; grey; n = 113 axons, 5 mice; Wilcoxon matched-pairs signed rank test). **(D)** Spontaneous (Sp.) and forepaw tactile-evoked (FP) Ca^2+^ peak amplitude in POm axons with evoked activity (n = 44 axons, 5 mice; Mann-Whitney test).

### The activity of POm axons in forepaw S1 during ‘action’ goal-directed behavior

Since POm axons within S1 increase their activity during forepaw stimulation, we next wanted to assess the activity of POm projection axons during tactile-based behavior. To test this, Ca^2+^ imaging of POm axons projecting to forepaw S1 was performed in mice (p50 - 70) trained in a goal-directed tactile detection task. Here, mice were trained to associate forepaw tactile stimulation (200 Hz, 500 ms) with a reward (see Methods; Figure 2A). If mice correctly responded by licking a reward port within 1.5 sec after receiving the tactile stimulus, a sucrose water reward (10 % sucrose water) was delivered. We refer to this behavioral paradigm as ‘action’ goal-directed task (action task). Mice rapidly learnt this task, taking on average 4.38 ± 0.37 days to reach expert level (> 80 % correct responses to tactile stimulation; Figure S3). Once expert, Ca^2+^ transients were recorded from POm axons within forepaw S1 while the mouse performed the action task (Figure 2B). Overall, POm axonal activity in forepaw S1 was significantly increased from the naïve state, with large Ca^2+^ transients evoked in 90 % of active POm axons during correct performance in the task (HIT; n = 105 of 113 axons, 11 mice; Figure 2C, D and Figure S4). To further assess the activity of these active POm axons, we categorized all axons according to their peak activity during baseline (−3 – 0 s pre stimulus), stimulus/response (response; 0 – 2 s post stimulus) or reward (2 – 4 s post stimulus) epochs of the tactile goal-directed behavior (Figure 2E). Here, POm axonal activity was greatest during the response epoch, increasing signaling by more than 4-fold above baseline (probability per trial, 0.08 ± 0.003 vs 0.32 ± 0.02; n = 418 axons, 11 mice; p < 0.0001; Figure 2F). POm axons also encoded reward information, with significantly increased activity above baseline during the reward epoch (probability per trial, 0.12 ± 0.007; n = 418 axons, 11 mice, p < 0.0225). However, when compared with the response-evoked activity, POm axons were significantly less active during reward, illustrating that POm axons preferentially encode behavioral response (p < 0.0001; Figure 2F). Direct comparison of the Ca^2+^ transient amplitudes from POm axons with both spontaneous and evoked activity (n = 239 axons, 11 mice) illustrates that Ca^2+^ transients evoked during the response epoch (0.972 ± 0.042 ΔF/F) were also significantly larger than transients evoked during both baseline (0.756 ± 0.02 ΔF/F, p = 0.0005) and reward delivery (0.679 ± 0.0219 ΔF/F; p < 0.0001), further highlighting the enhanced POm axonal signaling during the behavioral response (Figure 2G). Together, these results illustrate that POm increases the reliability, and content, of the information transferred to S1 during both response and reward delivery, with greatest activity during the behavioral response to tactile goal-directed behavior. Licking motion itself did not influence POm axon activity in forepaw S1, as there was an inverse relationship between licking frequency and POm axonal activity during the tactile goal-directed task (Figure 2H). Furthermore, spontaneous licking caused no detectable changes in the probability of Ca^2+^ transients above the spontaneous rate (0.09 ± 0.027; p = 0.29; n = 71 axons, 3 mice). Therefore, overall, POm axons in forepaw S1 encode behavioral response and, to a lesser extent, reward information during a tactile goal-directed task.

**Figure 2.**
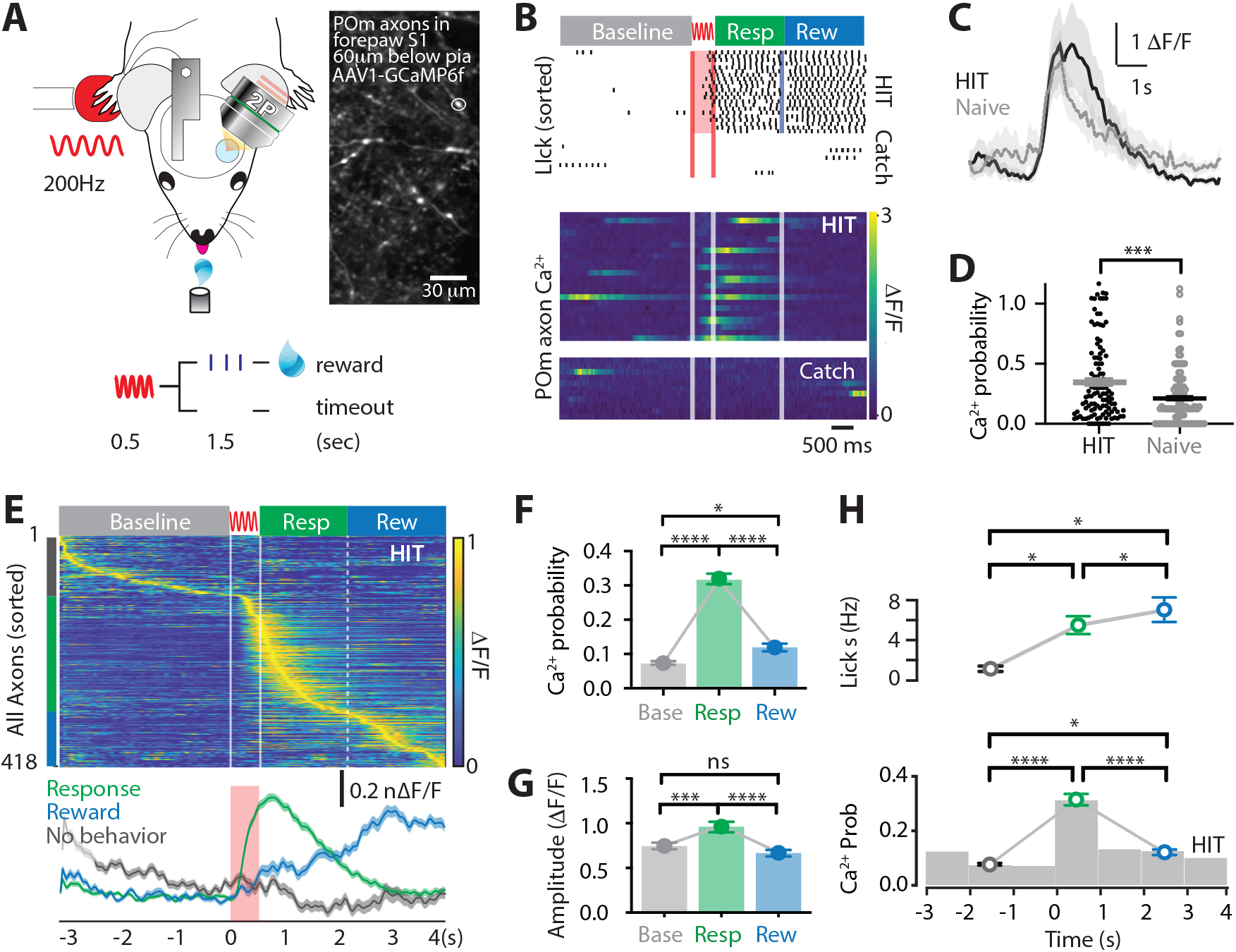
POm axonal projections in forepaw S1 have greatest activity during the behavioral response of a tactile goal-directed task. **(A)** Behavioral task design. Two-photon Ca^2+^ imaging of GCaMP6f-expressing POm axons in forepaw S1 was performed in head-restrained mice trained to report the detection of a tactile stimulus (200 Hz, 500 ms) by licking a reward port. Correct responses (HIT) were rewarded with sucrose water reward (10μl, 10% sucrose). **(B)** Top, Raster plot showing a typical behavioral response (licks) sorted into correct HIT performance and Catch (no-stimulus) trials. Grey, spontaneous; red, tactile stimulus; green, response epoch; blue, reward epoch. Blue line, reward delivery. Middle, Ca^2+^ activity pattern during correct performance and Catch trials from example POm axon in (A). **(C)** Mass average with SEM (shaded area) of all stimulus-evoked Ca^2+^ transients during correct goal-directed performance (HIT; black) and naïve (grey). **(D)** Probability of evoking a Ca^2+^ response during correct HIT behavior (black) compared with tactile-evoked activity in the naïve state (grey, n = 113 axons; Mann-Whitney test). **(E)** Top, Ca^2+^ activity pattern during HIT performance in the tactile goal-directed task sorted according to behavior during peak activity (grey, baseline; red, stimulus; green, response epoch; blue, reward epoch). Dashed line, reward delivery. Bottom, Average Ca^2+^ response in POm axons active during the stimulus and response epoch (green), reward epoch (blue); baseline (no behavior; grey). Red bar, stimulus delivery. **(F)** The probability of a Ca^2+^ transient in POm axons during baseline (grey), response epoch (green), reward epoch (blue). n = 418 axons, 11 mice. Friedman test + Dunn’s multiple comparisons test. **(G)** The amplitude of Ca^2+^ transients in POm axons evoked during baseline (grey), response epoch (green), reward epoch (blue). n = 239 axons, 11 mice with evoked Ca^2+^ transients. Friedman test + Dunn’s multiple comparisons test. **(H)** Top, Average lick frequency during spontaneous (grey), stim/response (green) and reward (blue) epochs during correct HIT behavior. Bottom, Histogram of Ca^2+^ transient probability in POm axons. * P < 0.05, *** P < 0.001, **** P < 0.0001.

### POm axon activity in forepaw S1 encodes correct tactile goal-directed behavior

We next assessed whether POm axonal activity in forepaw S1 changes according to behavioral performance. Upon receiving a tactile stimulus, mice had to lick a reward port within 1.5s to receive a sucrose water reward (HIT). However, if they did not respond during this epoch, then no water was delivered (MISS; Figure 3A). Despite performing at expert level, mice did not respond (MISS) to on average 12.63 ± 6.63 % of tactile stimuli. To assess whether POm axons also encode MISS behavior, evoked Ca^2+^ activity was directly compared in POm axons with both HIT and MISS activity (n = 159 axons, 6 mice). Compared to correct HIT behavior, POm axons in forepaw S1 were overall less active during MISS trials (Figure 3B). In addition to a decrease in the number of axons active during the tactile goal-directed task (by 51%), there was also a significant decrease in the probability of evoking an axonal Ca^2+^ event during the behavioral response in MISS trials (paired; HIT, 0.27 ± 0.02; MISS, 0.12 ± 0.01; p < 0.0001; Figure 3C). Behaviorally speaking, HIT and MISS trials differ in the mouse movement, which has been shown to increase overall brain activity (Stringer et al., 2019). To investigate whether the increased POm axonal activity during the HIT response to tactile goal-directed behavior is due to body movement, we compared the evoked Ca^2+^ activity in POm axons with catch trials where mice spontaneously licked for reward (false alarm). Despite false alarm trials having the same licking motion, POm axons were significantly less active than during HIT trials (probability per trial, 0.15 ± 0.01, n = 239 axons, 10 mice, p < 0.0001; Figure 3C). Therefore, this data further suggests that POm axonal activity is not simply due to licking behavior. There was also a significant decrease in the probability of evoking an axonal Ca^2+^ event during the reward epoch in MISS trials (HIT, 0.13 ± 0.02; MISS, 0.09 ± 0.01; p = 0.0408; n = 159 axons, 6 mice; Figure 3D). During MISS trials, POm axonal activity was similar to baseline rates (0.07 ± 0.007; p > 0.999; n = 159 axons, 6 mice; Figure 3D). Together, this data suggests that the POm encodes behavioral performance, increasing the information transfer between the POm and forepaw S1 during correct HIT behavior during both the response and reward epochs in a tactile goal-directed task.

**Figure 3.**
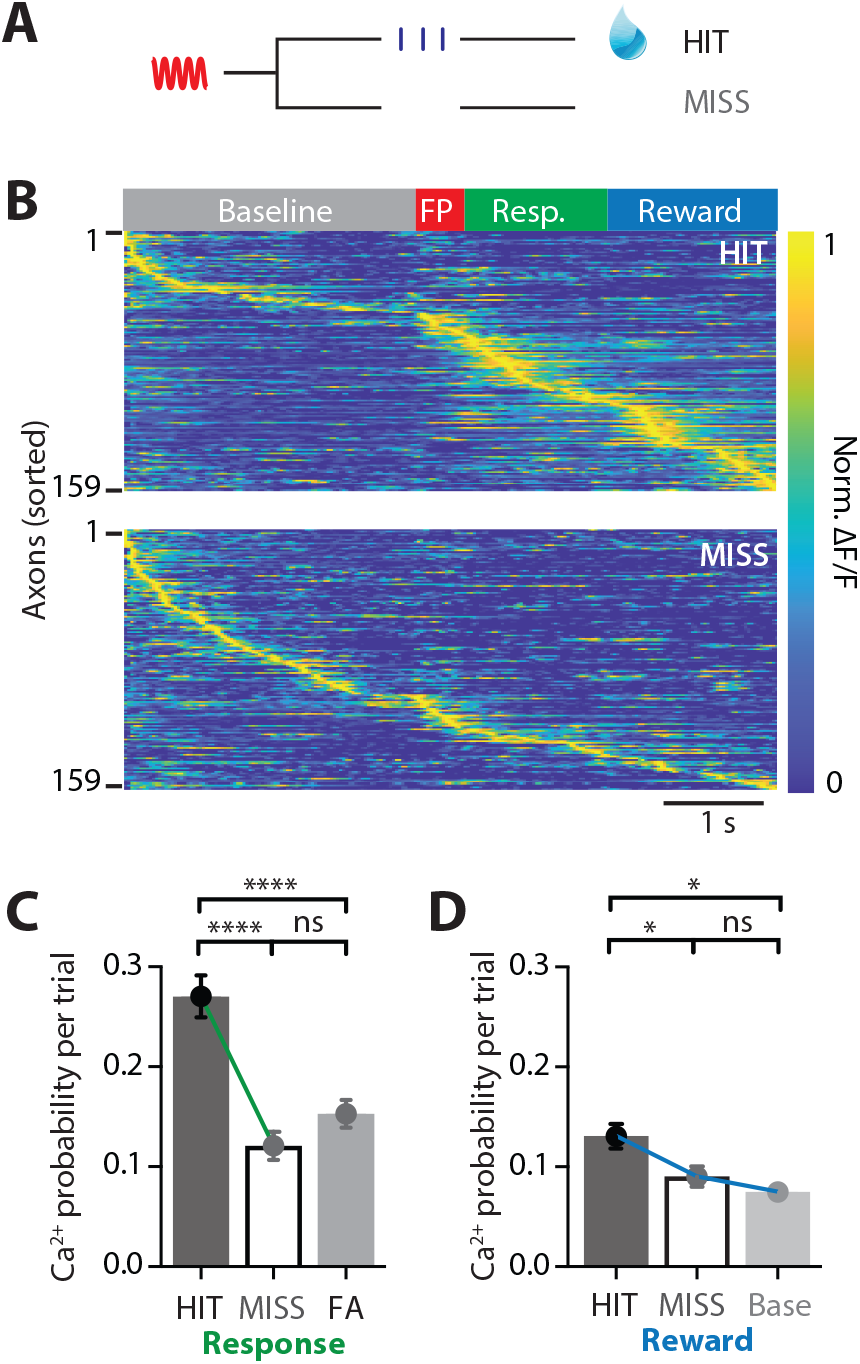
POm axonal projections in forepaw S1 have greatest activity during the behavioral response of a tactile goal-directed task. **(A)** Behavioral task design. Two-photon Ca^2+^ imaging of GCaMP6f-expressing POm axons in forepaw S1 was performed in head-restrained mice trained to report the detection of a tactile stimulus (200 Hz, 500 ms) by licking a reward port. Mice received sucrose water reward (10μl, 10% sucrose) during correct responses (HIT), whereas incorrect responses (MISS) were unrewarded. **(B)** Ca^2+^ activity patterns in POm axons with Ca^2+^ transients evoked during HIT (top) and MISS (bottom) behavior during the tactile goal-directed task (n = 159 axons, 6 mice). Grey, baseline; Red, stimulus; Green, response epoch; Blue, reward epoch. Each row is an axon normalized to maximum fluorescence and sorted by the timing of the peak amplitude. **(C)** The probability of a Ca^2+^ transient evoked during the response epoch in HIT (solid), MISS (empty) and false alarm (FA; dark grey) behavior. Wilcoxon matched-pairs signed rank test (HIT vs MISS) and Mann-Whitney test (FA vs HIT and MISS) **(D)** The probability of a Ca^2+^ transient evoked during the reward epoch in HIT (solid), MISS (empty) and baseline (light grey). Friedman test + Dunn’s multiple comparisons test. * P < 0.05, *** P < 0.001, **** P < 0.0001.

### POm axon activity in forepaw S1 during suppression of a goal-directed action

Goal-directed behavior requires motor actions to be suppressed once they are no longer appropriate to achieve the current goal (Jahanshahi et al., 2015). To investigate the involvement of the higher order thalamus during suppression of a previously learned goal-directed action, we performed Ca^2+^ imaging from POm axons during a modified goal-directed paradigm. Here, mice previously injected with the Ca^2+^ indicator GCaMP6f in the POm were trained in the ‘action’ goal-directed task (as above). Once expert (> 80 % correct responses to tactile stimulation), the behavioral paradigm was changed such that the mice only received the reward if they suppressed licking in response to the tactile stimulus (Figure 4A). We refer to this behavioral paradigm as ‘action-suppression’ goal-directed task (suppression task). To monitor cognitive engagement, dynamic changes in pupil diameter were recorded while mice were performing the suppression task. Despite the enforced behavioral (licking) suppression, mice were highly engaged in the task. When compared with the action goal-directed task, there was no significant difference in peak pupil diameter during baseline (0.32 ± 0.05 vs 0.29 ± 0.06 mm, p = 0.3121), pre-stimulus (0.35 ± 0.06 vs 0.31 ± 0.07 mm, p = 0.2193), and post-stimulus (0.44 ± 0.07 vs 0.41 ± 0.09 mm; p = 0.6872; n = 6 mice; Figure 4B). Furthermore, there was no significant difference in correct performance rates during the action (83 ± 5 % correct; n = 11 mice) and suppression (86 ± 7 %; n = 6 mice; p = 0.57) tasks. POm projections in S1 were highly active during the suppression task, with evoked Ca^2+^ transients that were significantly larger in amplitude than spontaneous activity (0.99 ± 0.06 ΔF/F vs 1.19 ± 0.06 ΔF/F; n = 144 axons, 6 mice; p = 0.0002). Similar to the action task, most POm axons were maximally active during the response epoch in correct HIT trials (0.24 ± 0.03; n = 144 axons, 6 mice; Figure 4C). Likewise, a subset of axons were also active during the reward epoch of the suppression task, significantly increasing activity above baseline (probability per trial, 0.09 ± 0.01 vs 0.18 ± 0.02; n = 95 axons, 6 mice; p <0.0001; Figure 4C). To assess whether POm activity was also decreased during MISS behavior in the suppression task, evoked Ca^2+^ activity was directly compared in POm axons during the during the response epoch. Similar to the action task, POm axons in forepaw S1 were less active during MISS behavior, with a significant decrease in the probability of response-evoked activity in MISS trials compared to HIT trials (HIT, 0.24 ± 0.03 vs MISS, 0.15 ± 0.03; n = 144/53 axons, 6 mice; p = 0.029; Figure 4D). Here, MISS behavior involves incorrectly licking for reward during the response epoch further illustrating that POm axonal activity in S1 does not signal licking behavior. Taken together, in both the action and suppression tasks, POm axons in forepaw S1 encode behavioral response during correct performance (HIT trials). On average, the peak amplitudes (1.19 ± 0.06 vs 1.26 ± 0.05 ΔF/F, p = 0.3812) and durations (623 ± 50 vs 666 ± 35 ms; p = 0.2234) of Ca^2+^ transients evoked during the action and suppression tasks were comparable (Figure 4E). However, during the suppression task, the probability of evoked POm signaling during the response epoch was significantly decreased compared to the action task (p = 0.0007; Figure 4F). This is in contrast to the similar probability of evoked POm signaling during the reward epoch (p = 0.87; Figure 4F). Together, these results further support the encoding of correct goal-directed active behavior by POm axons within forepaw S1.

**Figure 4.**
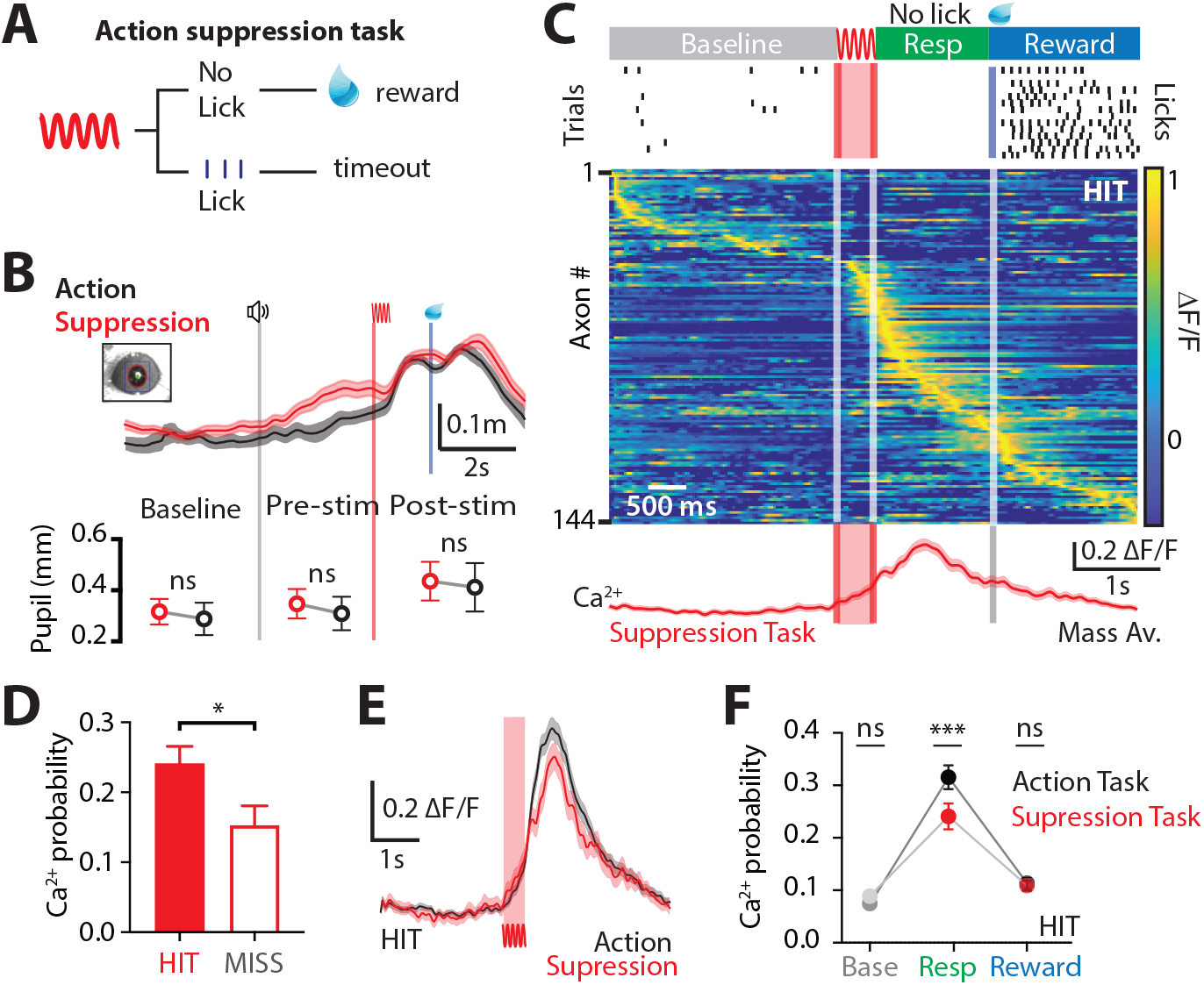
Ca^2+^ dynamics in POm axonal terminals during suppression of a goal-directed action. **(A)** Behavioral task design. Two-photon Ca^2+^ imaging of POm axon terminals was performed in head-restrained mice trained to suppress a previously learned goal-directed action. Mice were trained to withhold licking in response to forepaw stimulation (200 Hz, 500 ms) for 1.5 seconds to get a reward (10 μl, 10 % sucrose water). **(B)** Top, Average pupil diameter with SEM (shaded area) during correct performance at the ‘suppression’ goal-directed task (red) and ‘action’ goal-directed task (black). Bottom, Comparison of pupil dilation during the ‘action’ and ‘suppression’ goal-directed tasks in baseline, pre-stim and post-stim epochs (n = 6 mice: Wilcoxon matched-pairs signed rank test). Grey line, trial start; red line, stimulus, blue line, reward delivery. **(C)** Top, Raster plot showing the typical licking response during correct performance of the task. Grey, spontaneous; Red, stimulus; Green, response epoch; Blue, reward epoch. Blue line, reward delivery. Middle, Normalized Ca^2+^ activity pattern during correct performance of the suppression task (n = 144 axons, 6 mice). Each row is an axon normalized to maximum fluorescence and sorted by the timing of the peak amplitude. Bottom, Mass average with SEM (shaded area) of the normalized Ca^2+^ activity pattern during correct performance. **(D)** Probability of evoking a Ca^2+^ transient during HIT (correct suppression of licking behavior; red) and MISS (no suppression of licking behavior; red empty). n = 144 axons, 6 mice; Wilcoxon matched-pairs signed rank test. **(E)** Overlay of the mass average with SEM (shaded area) of the normalized Ca^2+^ activity pattern during correct performance in the Suppression goal-directed task shown in (C) (red) and Action goal-directed task (black). **(F)** Probability of evoked Ca^2+^ transients during baseline, response, and reward epochs in the ‘suppression’ goal-directed task (red) and ‘action’ goal-directed task (black). Mann-Whitney test. ** P < 0.01, *** P < 0.001, **** P < 0.0001.

### The influence of POm input during goal-directed behavior

How crucial is POm signaling during active goal-directed behavior? To test this, the inhibitory opsin, archaerhodopsin (ArchT; AAV1.CAG.ArchT.GFP.WPRE.SV40, 60 nl), was unilaterally injected into the POm. The effectiveness of 565 nm LED photo-inhibition of POm neurons expressing ArchT was tested using patch clamp electrophysiology in the thalamic brain-slice preparation. Here, although photo-inhibition did not completely abolish action potentials in POm neurons, the evoked firing rate was significantly decreased by 64 ± 13 % (p = 0.031; n = 6 neurons; Figure S5). To test the influence of this decrease in POm activity on active goal-directed behavior, a fiber-optic cannula was chronically inserted into the POm which was previously injected with ArchT (see methods and Figure 5A). Mice were trained in the ‘action’ goal-directed task, with the duration of training and baseline performance not affected by the injection (Figure S6). Once expert (> 80 % correct performance), the POm was photo-inactivated with interleaved yellow LED light (565 nm, 5 mW, 2 s) during the response epoch (see Methods, Figure 5B). Here, partial photo-inactivation of the POm produced a significant reduction in the overall behavioral performance (d prime, 2.58 ± 0.15 vs. 2.23 ± 0.26; n = 9 mice; p = 0.04, Figure 5C) while no change was observed in the control group injected with GFP in the POm (d prime, 2.62 ± 0.24 vs. 2.83 ± 0.29; n = 9 mice; p = 0.26; Figure 5D). Furthermore, POm photo-inactivation did not alter licking behavior as there was no significant difference in the latency to the first lick (control, 351 ± 29 ms vs ArchT, 342 ± 26 ms, n = 9 mice, p = 0.1282, Figure 5E). Taken together, decreasing POm activity produced a small effect on correct performance in a goal-directed task, illustrating a direct, but not driving, influence of the higher-order thalamus in goal-directed behavior.

**Figure 5.**
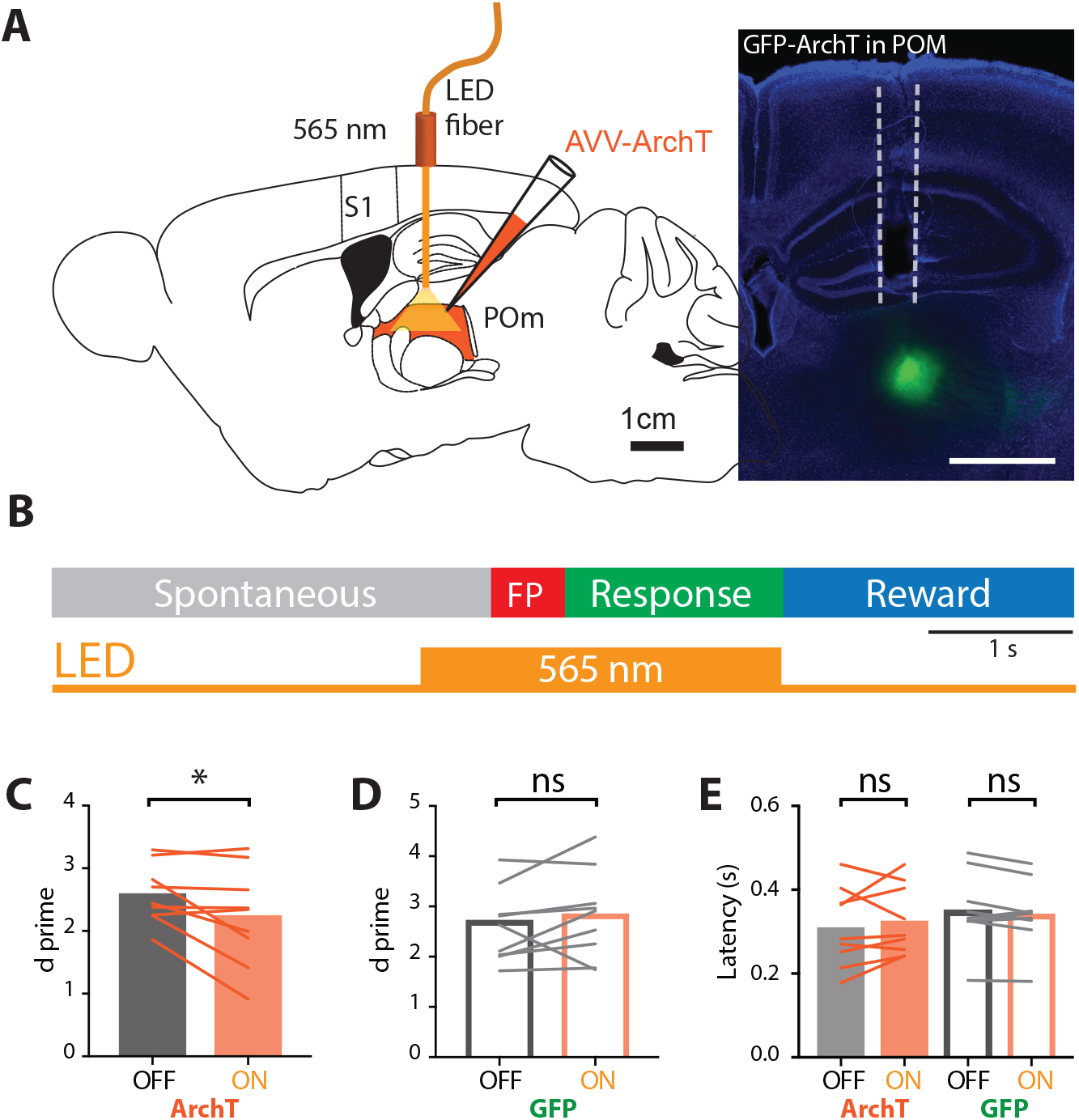
Optogenetic inactivation of the POm during the Action goal-directed task. **(A)** Left, Experimental design. The inhibitory opsin, archaerhodopsin (ArchT) was unilaterally injected into the POm and a fiber-optic cannula was chronically inserted into the brain. Right, Localized ArchT spread in POm and fiber optic track (dotted line), bar = 1 mm. **(B)** The POm was photo-inactivated (590 nm, 5 mW, 2 s) during the stimulus (FP, red) and response (green) epochs in mice expert in the ‘action’ goal-directed task. **(C)** Behavioral performance for light OFF versus ON trials in mice injected with ArchT (n = 9 mice). **(D)** Behavioral performance for light OFF versus ON trials in control mice injected with GFP (n = 9 mice). **(E)** Latency to the first response lick in mice injected with ArchT (sold) and control GFP (empty). Individual values are shown. Wilcoxon matched-pairs signed rank test *P < 0.05.

### Tactile-evoked responses in POm neurons in the awake state

Since the POm contributes to the correct performance of the tactile goal-directed task, we next asked what influence does the POm have on pyramidal neuron activity in the somatosensory cortex. In particular, we investigated the influence of POm input on cortical L2/3 pyramidal neurons, as they send and receive long-range projections from other cortical areas (Oh et al., 2014; Yamashita et al., 2018), making them the perfect hub for the integration of cortical and sub-cortical information. To address this, we needed to first establish the action potential response to tactile stimulus in POm neurons. Juxtacelluar recordings were performed from POm neurons in awake mice at an average recording depth, 2926 ± 38 μm (see methods, Figure 6A and Figure S7). During forepaw stimulus (500 ms; 200 Hz), POm neurons evoked a variable number of action potentials (Figure 6B) with an average latency to the first action potential of 95.9 ± 8.2 ms (289 trials, n = 18 neurons; 4 mice). The firing rate evoked during tactile stimulus was greater than the spontaneous rate, with a significant decrease in inter-spike interval compared to spontaneous action potentials (spont, 115.1 ± 9.61 ms; FP, 106 ± 16.1 ms; p < 0.0001; n = 289 trials, 18 neurons, 4 mice; Figure 6C).

**Figure 6.**
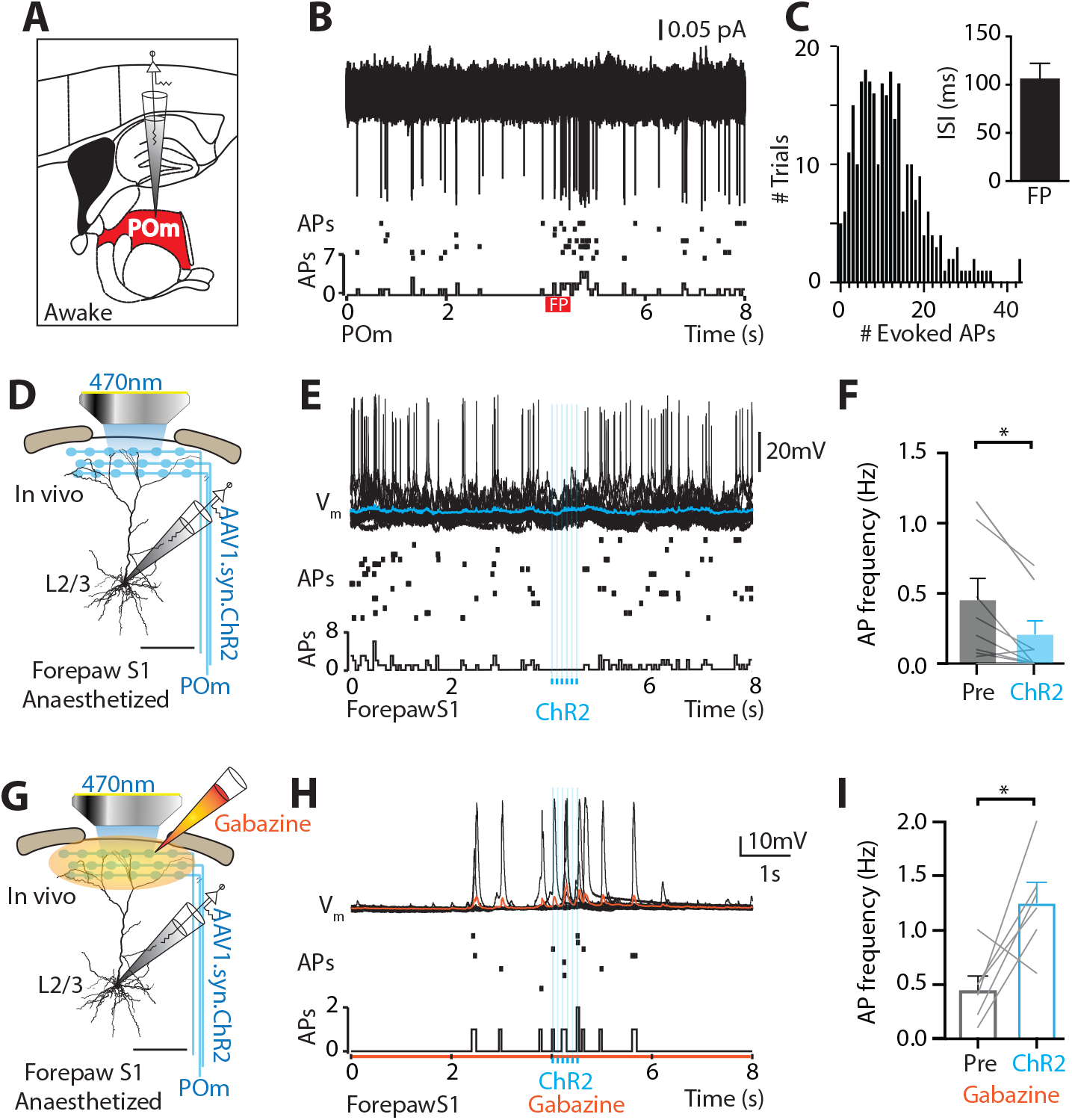
POm fires sustained action potentials during tactile stimulus which, when POm projection axons are optogenetically photoactivated, causes inhibition in cortical L2/3 pyramidal neurons. **(A)** Experimental design. Juxtacellular voltage recordings were performed from POm neurons in awake mice. **(B)** Top, Overlay of voltage responses from a typical POm neuron during tactile stimulus to the contralateral forepaw (200 Hz; 500 ms). Middle, Raster of action potentials. Bottom, Histogram of action potentials. **(C)** Histogram of evoked action potentials in POm neurons in response to tactile forepaw stimulus for all recorded neurons (n = 18 neurons, 4 mice). Inset, Action potential interspike interval (ISI) for all spontaneous (grey) and tactile-evoked (black) action potentials in POm neurons (n = 289 trials, 18 neurons, 4 mice; p < 0.0001; Wilcoxon matched-pairs signed rank test). **(D)** Experimental design. *In vivo* whole-cell patch clamp recordings were performed from L2/3 pyramidal neurons in forepaw S1 during sustained photoactivation (470 nm) of ChR2-expressing POm axons. **(E)** Top, Overlay of voltage responses from a typical L2/3 pyramidal neuron evoked by sustained (6x; 10 ms; 90 ISI) photoactivation of POm projection axons in forepaw S1. Average, blue. Middle, Raster of action potentials. Bottom, Histogram of action potentials. **(F)** Evoked action potential frequency pre- (grey) and during (ChR2; solid blue) sustained photoactivation of POm axons within forepaw S1 (n = 10 neurons, 9 mice; Wilcoxon matched-pairs signed rank test). **(G)** Experimental design. *In vivo* whole-cell patch clamp recordings were performed from L2/3 pyramidal neurons in forepaw S1 during sustained photoactivation (460 nm) of ChR2-expressing POm axons and cortical application of Gabazine. **(H)** Top, Overlay of voltage responses from a typical L2/3 pyramidal neuron evoked by sustained (6x; 10ms; 90 ISI) photoactivation of POm projection axons in forepaw S1 during cortical application of Gabazine. Average, blue. Middle, Raster of action potentials. Bottom, Histogram of action potentials. **(I)** Evoked action potential frequency pre- (grey) and during (ChR2; empty blue) sustained photoactivation of POm axons within forepaw S1 during cortical application of Gabazine (n = 6 neurons, 5 mice; Wilcoxon matched-pairs signed rank test). *P < 0.05, ****P < 0.0001.

### The influence of sustained POm input on layer 2/3 pyramidal neurons in forepaw S1

We next investigated the influence of this evoked POm input on the firing activity of cortical neurons. Patch clamp recordings were performed from L2/3 pyramidal neurons in the forepaw S1 in urethane (1.3g/Kg) anaesthetized mice which were previously injected with the excitatory opsin, ChannelRhodopsin2, in the POm (see methods; Figure 6D, S1 and S2). Based on the evoked firing rate recorded from POm neurons during forepaw stimulus (Figure 6C), POm axons were photo-activated with a train of brief LED pulses (10 ms; ISI, 90 ms; 470 nm) for the equivalent duration of the forepaw stimulus (500 ms; Figure 6E and S8). Using this stimulus paradigm, photo-activation of sustained POm input caused a significant decrease in the action potential firing rate in L2/3 pyramidal neurons (0.455 ± 0.150 vs 0.210 ± 0.094 Hz; p = 0.008; n = 10 neurons, 9 mice; Figure 6F). Considering the known cortical inhibitory microcircuits activated by POm (Audette et al., 2017), we next tested whether the inhibition of L2/3 pyramidal neurons by POm input was mediated by GABA_A_-receptor activation. The GABA_A_-receptor agonist Gabazine (10 μM) was applied onto the cortical surface during sustained photoactivation of POm axons in forepaw S1 (Figure 6G). Blocking GABA_A_-receptors abolished the POm-evoked inhibition of L2/3 pyramidal neuron firing, with sustained POm input instead resulting in a significant increase in action potentials (pre, 0.45 ± 0.13 vs ChR2, 1.26 ± 0.19 Hz; p = 0.0316; n = 6 neurons, 5 mice; Figure 6H and I). These results illustrate that sustained POm input to forepaw S1 results in GABA_A_-mediated inhibition of cortical layer 2/3 pyramidal neurons activity.

## DISCUSSION

The results presented here highlight the role of the POm during sensory-based goal directed behavior. We used electrophysiology and two-photon Ca^2+^ imaging to illustrate that POm axons encode tactile stimulus in the naïve state, increasing action potential generation throughout tactile stimulation of the forepaw. During tactile goal-directed behavior, POm axons in forepaw S1 further increased activity during the response and, to a lesser extent, reward epochs. Therefore, the POm does not simply code sensory stimulus, but rather plays an important role during the behavioral response in a goal-directed task. Despite the known importance of the thalamocortical circuit in brain function, to our knowledge, this is the first study which directly records thalamic axonal activity in the cortex during a goal-directed behavior.

Since POm receives inputs from M1 (Yamawaki and Shepherd, 2015), it could be speculated that the increased axonal activity during the behavioral response has a motor origin. In fact, previous studies have shown that the thalamus is a circuit hub in motor preparation (Guo et al., 2017). However, while a general increase in POm activity has been reported during active states (Urbain et al., 2015), encoding of whisking related movement in the POm is relatively poor (Moore et al., 2015). Our findings illustrate that POm input in forepaw S1 does not specifically encode movement, as 1) POm activity is enhanced during the response epoch in both the action (licking) and suppression (no licking) tasks, 2) there is not a strong correlation between POm activity and licking frequency, and 3) POm activity is minimal during spontaneous licking and false alarm trials. Here, we illustrate that POm activity instead reflects task performance with greater signaling in forepaw S1 during correct HIT behavior in both the action and suppression task. Increased activity in the higher order thalamus has been associated with the expected value and significance of rewarded sensory stimuli (Komura et al., 2001) and may reflect learned-dependent strengthening of specific POm thalamocortical synapses (Audette et al., 2019). Indeed, our findings show that POm activity is enhanced during reward delivery in the tactile goal-directed task, although the absolute POm signaling is less than during the behavioral response.

In both the anaesthetized (Mease et al., 2016) and awake (Urbain et al., 2015) preparations, POm neurons increase their firing rate, generating action potentials in response to sensory input. Using direct juxtacellular cell-attached recordings, we showed that POm neurons typically fire a train of action potentials in response to tactile stimulus, at similar rates to whisking (Urbain et al., 2015). We investigated the influence of this sustained POm activity on the cortex using the precise control afforded by optogenetics. In particular, we investigated the influence of POm input on cortical L2/3 pyramidal neurons, as they send and receive long-range projections from other cortical areas (Oh et al., 2014; Yamashita et al., 2018), making them an ideal hub for cortical information processing. Furthermore, since they effectively alter the gain of layer 5 pyramidal neurons (Quiquempoix et al., 2018), any changes in L2/3 pyramidal neuron firing will have downstream effects on cortical output. During sustained photoactivation of POm axons in forepaw S1, action potentials in L2/3 pyramidal neurons were strongly inhibited. This contrasts with enhanced cortical responses during a single brief photo-activation of POm input (Gambino et al., 2014; Mease et al., 2016; Zhang and Bruno, 2019). The opposing influence of POm input on cortical neurons highlights the complexity of the higher order thalamocortical microcircuit, with POm input directly targeting both pyramidal neurons and various subtypes of interneurons (Audette et al., 2017). Our findings suggest that during goal-directed behavior, L2/3 pyramidal neurons in forepaw S1 would be suppressed by input from the higher order thalamus. In line with these findings, layer specific cortical interneurons have been previously reported to be recruited by thalamic input during sensory behavior (Yu et al., 2019). Further studies are required to delve into the influence of the higher order thalamus on the cortex during different sensory inputs and behaviors.

During partial photo-inhibition of POm neurons, behavioral performance in tactile goal-directed behavior was disrupted. The behavioral effect was small (but robust) which may be due to the following. Firstly, we illustrate that photo-inhibiting LED light (565 nm) caused a significant decrease in the evoked action potential rate in POm neurons expressing Archaerhodopsin. However, firing activity was not abolished therefore during the goal-directed behavior, the POm was still active, albeit at a third of the evoked firing rate. Secondly, previous studies have illustrated that similar sensory-based goal-directed behaviors do not require primary cortical areas (Hong et al., 2018). Therefore, it is not expected that partially inhibiting an input stream to the forepaw S1 would have large effect on the behavioral performance. Combined with the reported increase in POm axonal activity during correct performance, this further supports the finding that POm encodes correct goal-directed action.

Considering that patterns of cortical activity during behavior have been associated with task engagement, brain state, attention, motivation or reward (Kobak et al., 2016; Lacefield et al., 2019; Poort et al., 2015; Poulet and Petersen, 2008; Reimer et al., 2014), we monitored pupil dynamics during the goal-direct task. We report that while overall POm activity increased concomitantly with pupil diameter during the behavioral response, this trend was reversed during reward delivery. However, by sorting POm axons according to their peak activity during the tactile goal-directed task, we revealed a subgroup of POm axons highly responsive during the reward epoch. This finding highlights the heterogeneity of the higher order thalamus, with subsets of POm axonal projections specifically encoding either the behavioral response or reward delivery. Overall, however, this study illustrates that the POm has greatest signaling during the response to a tactile goal-directed task.

In summary, we show that the higher order thalamus plays an active role during cognitive functions such as goal-directed behavior. This finding expands the known roles of the higher-thalamic nuclei, from sensory encoding to action selection. Overall, the thalamus is not a simple relay system. It encodes and drives goal-directed behaviors which are crucial for survival in a dynamic environment.

## Acknowledgments

We would like to thank the Palmer laboratory and Matthew Larkum for their helpful discussions and comments on the manuscript. We would also like to thank Verena Wimmer for her POm expertise and Ronny Bergmann, Viktor Bahr, Jens Kremkow and Robert Sachev for use of their pupil-tracking software.

## Author contributions

D.L and L.M.P designed the experiments. D.L performed and analyzed all Ca^2+^ imaging and optogenetic experiments. S.P. performed and analyzed the *in vivo* POm recordings. M.R. performed and analyzed the *in vitro* POm recordings. D.L and L.M.P wrote the manuscript.

## Data and materials availability

All data is available in the main text or in the Supplementary material.

## Online Methods

All procedures were approved by the Florey Institute of Neuroscience and Mental Health Animal Care and Ethics Committee and followed the guidelines of the Australian Code of Practice for the Care and Use of Animals for Scientific Purposes.

### Mice

Wild type C57BL/6 female mice (PN30 - PN80) were used in this study. Mice were housed in groups of six in a 12:12 natural light/dark cycle. All behavioral tests were performed during the light phase.

### Virus injection

All surgical procedures were conducted under isoflurane anesthesia (~1-2% in O2). Body temperature was maintained at ~36 °C and the depth of anesthesia was monitored throughout the experiment. Mice (~PN30 - 40) were placed in a stereotaxic frame (Narishige) and eye ointment was applied to the eye to prevent dehydration. The skin was disinfected with ethanol 70% and betadine before lidocaine (1%, wt/vol) was topically applied to the wound edges for additional local anesthesia. An incision in the skin (10 mm) was made to expose the skull and a small craniotomy (~0.5 × 0.5 mm) was made over the left posteromedial (POm) complex of the thalamus using the following stereotaxic coordinates: rostrocaudal (RC), −1.7 mm; mediolateral (ML), −1.25 mm; dorsoventral (DV), −3.00 mm from bregma. Virus (60 nl, AAV1.Syn.GCaMP6f.WPRE.SV40; AAV1.CAG.ArchT.GFP.WPRE.SV40; or AAV1.hSyn.ChR2(H134R)-eYFP.WPRE.hGH) was slowly injected from a glass pipette (Wiretrol, Drummond) for at least 5 min using an oil hydraulic manipulator system (MMO-220A, Narishige). The skin was then sutured and Meloxicam (3mg / Kg) was injected intraperitoneally (i.p.) for additional post-operative analgesia and anti-inflammatory action. Mice were then returned to their home cage for recovery.

### Chronic cranial window surgery

Mice previously injected with the Ca^2+^ indicator GCaMP6f were anaesthetized (isoflurane, ~1-2 % in O2, vol/vol) and body temperature was maintained at ~36 °C and the depth of anesthesia was monitored throughout the experiment. Eye ointment was applied to prevent dehydration and the top of the head was disinfected with ethanol 70 % and betadine and lidocaine (1 %, wt/vol) was topically applied for additional local anesthesia. The skin covering the skull was removed, and a craniotomy was performed over the left forepaw area of the primary somatosensory cortex (centered at coordinates: RC, 0 mm; ML, −2.3 mm; from bregma). The dura was left intact and a circular coverslip (3 mm diameter) was placed over the open craniotomy and seal-attached to the skull with acrylic glue. A custom-made aluminum head bar (2 × 1 × 0.1 cm) was then attached to the skull for head-fixation using dental cement (C&B metabond, Parkell Inc). Meloxicam (3 mg / Kg) was injected i.p. for additional post-operative analgesia and anti-inflammatory action. Mice were then returned to their cages to recover until behavioral training (~ 2 weeks).

### Habituation and behavior

Mice were trained to perform a goal-directed tactile task using a custom made behavioral platform (Micallef et al., 2017). A three to four day habituation period preceded the beginning of the operant conditioning. During this period mice were handled and acclimatized to the behavioral setup. Mice were head restrained for incremental periods of time until habituated to head restraint. To maximize task engagement, a day prior to the beginning of behavioral training, mice were water restricted (1 ml/day of 10% sucrose water) and from this day onward this water regimen was maintained until the end of the experiment. Behavioral sessions lasted ~ 300 trials during which the mice typically obtained their daily water intake (1 ml per day) otherwise extra water was supplemented. Ca^2+^ imaging was performed following this habituation phase for naïve data.

### Behavioral platform

Mice were head-fixed to the recording frame and their paws rested unaided on either an active (contralateral) or inactive (ipsilateral) rod coupled to a stepper motor driven by an Arduino Uno microprocessor. The stepper motor delivered a pure frequency forepaw tactile stimulus (500 ms, 200 Hz). A water port was used to deliver a water reward (10ul, 10% sucrose water) and licking frequency was recorded via a custom-made piezo-based lick sensor attached to the lick port. All behavioral tests were carried out in the dark while the animal behavior was monitored with an infrared sensitive camera (Microsoft lifecam). During the first training sessions, mice were habituated to tactile stimulus and reward delivery (typically 1-2 sessions). To establish an association between stimulus and reward, mice were able to self-initiate a trial by licking the water port which instantaneously triggered both stimulus and reward. After this habituation phase, operant conditioning was performed. **Action goal-directed task**: Background white noise (~40 DB) was played for the duration of each trial to indicate task onset and mask non-task related sounds. Tactile stimulation (200 Hz, 500 ms) was delivered after a 3-sec baseline period. Following stimulus presentation, mice were given a 1.5 sec interval to report the detection of the tactile stimulus by licking the lickport (response epoch), after which reward was made available and cued by an auditory sound (400 Hz, 200 ms). Mice were then given a 2-sec time window to retrieve the reward after which the trial terminated followed by an inter trial interval (ITI) of randomized duration (between 4 – 7 sec). Only correct responses (licks during the response epoch) were rewarded (Correct) while failure to report stimulus detection was considered an incorrect response (Incorrect). Trials with no tactile stimulation (catch trials) were randomly interleaved with stimulus trials. Licking within the response epoch during a catch trial was considered a false alarm (FA) and punished with a timeout of incremental duration (2 - 7 sec) while withhold licking was the correct response which was not rewarded, correct rejection (CR). Implementing catch trials and randomized ITI ensured that animals could not solve the task by adopting a time-based strategy. To facilitate learning, during the first training session the frequency of stimulus/catch trials was set to 90 / 10 %, respectively. The frequency of catch trials was progressively increased up to 40 % and maintained at this ratio until mice could reliably perform at expert level (≥ 80 correct response rate). On average, mice reached expert level within 4.38 ± 0.37 training sessions. **Action-suppression goal-directed task**: Background white noise (~40 DB) was played for the duration of each trial to indicate task onset and mask non-task related sounds. As in the action goal-directed task, tactile stimulation (200 Hz, 500 ms) was delivered after a 3-sec baseline period. However, following stimulus presentation, mice were trained to withhold their licking for a 1.5-sec interval. Mice were then given a 2-sec time window to retrieve the reward after which the trial terminated followed by an inter trial interval (ITI) of randomized duration (between 4 – 7 sec). Correct suppression of licking during this epoch was rewarded with sucrose water (10 μl, 10 %). Conversely, if mice licked during this interval (early lick) no reward was delivered and the trial was aborted. Catch trials were used as in the action goal-directed task. Mice learned to reliably suppress licking (≥ 80 % correct response rate) after an average 6 ± 0.85 training sessions (Supplementary Fig. 6).

### Two photon Ca^2+^ imaging

Imaging of POm axons in forepaw S1 expressing the Ca^2+^ indicator GCaMP6f was performed in awake behaving mice through a chronic cranial window approximately 3 weeks after virus injection. Head-fixed mice were placed under a two-photon microscope (Thorlabs A-scope) and POm axons located 48 ± 6.8 μm below the pia surface were excited using a Ti:Sapphire laser (SpectraPhysics MaiTai Deepsee) tuned to λ = 940 nm and passed through a 16x water immersion objective (Nikon, 0.8 NA). GaAsP photomultiplier tubes (Hamamatsu) were used for detection. The field of view spanned 512 × 512 pixels and images were acquired at 30 Hz. To minimize photo-damage, the excitation power was adjusted online to the minimal value sufficient to record Ca^2+^ transients and the number of imaged trials for a given field of view (FOV) was restricted to a maximum of 40. During each trial, animal behavior was monitored with an infrared sensitive camera (Microsoft Lifecam). Forepaw position on the tactile stimulator was analyzed post-hoc and any trials where the forepaw was not in contact with the stimulation apparatus were removed from further analysis.

### Cannula implant and photo-inactivation of POm complex

For optical inactivation of the POm complex, mice were injected ipsilaterally into the left POm with the inhibitory opsin AAV1.CAG.ArchT.GFP.WPRE.SV40 (60 nl). Following virus injection, a custom-made fiber optic cannula (FT400EMT, 400 um 0.39 NA, 2.5 mm fiber, Thorlabs) was slowly lowered down the injection track using a stereotaxic arm until the desired depth was reached (2.5 mm from pia). Dental cement (C&B metabond, Parkell Inc) was then applied around the edges of the cannula to secure it to the skull and left to dry for ~ 5 minutes. The same dental cement was used to attach a custom-made aluminum head bar (2 × 1 × 0.1 cm) to the skull for head-fixation. Meloxicam (3 mg/Kg) was injected i.p. for additional post-operative analgesia and anti-inflammatory action. Mice were then returned to their cages to recover until behavioral training (~ 3 weeks). **Behavioral procedures:** After recovery, mice were trained on the Action goal-direct task (see Habituation and Behavior). All behavioral procedures were performed using the Bpod behavioral platform (Bpod State Machine r1, Sanworks). Once mice reached expert level (≥ 80% correct response rate), an optogenetic experimental session was performed. Photo-inactivation of the POm complex was achieved by delivering a light pulse (565 nm, 2-sec, 5 mW) through a 400 μm optical fiber (FT400EMT, Thorlabs) directly inserted into the cannula (FT400EMT, 400 um 0.39 NA, 2.5 mm fiber, Thorlabs). The light pulse was delivered 500 ms prior to stimulus onset to ensure inactivation of the POm complex. A LED light source (LEDD1B, Thorlabs) coupled to a 565 nm LED filter (M565F3, Thorlabs) was used to generate the photo-stimulus. A custom-made light shield was placed over the animal’s head to prevent scattered light from entering the animal visual field. During a typical experimental session (~300 trials), LED-ON and LED-OFF trials were randomly interleaved at a rate of 50% each. Custom routines in Matlab were used to operate the behavioral platform and data acquisition. For control experiments, mice were stereotaxically injected into their left POm (see virus injections) with AAV1-PAM MuseeGFP (60 nl) and experiments carried out as above.

### Pupil tracking and analysis

To monitor engagement during the task, pupil tracking was performed in a subset of mice previously trained on the Action goal-directed task for the ArchT experiments (see above). Pupil tracking was performed when mice were expert on both the Action task and the Action-Suppression task (see Habituation and Behavior). Pupil tracking was also performed during the transition between these tasks (switching) when their correct response rate dropped to chance level (~50 %). Mice were head-fixed and the right eye illuminated with infrared light (850 nm LED, Thorlabs). This illumination did not affect pupil diameter. Behavioral sessions were performed on the same apparatus used for two photon imaging inside an aluminum soundproof optical enclosure. However, some illumination (3.48 lux) was provided as we found that the pupil became maximally dilated and a-dynamic in complete darkness. An IR sensitive camera (Basler aCA1300-200 um) mounting a 50 mm lens (Kowa 50mm / F2.8) was used to image pupil dynamics at 15 frames per second. Frames were triggered externally using an Arduino microprocessor connected to a Bpod (Bpod State Machine r1, Sanworks) which was then used to operate the behavioral paradigm. Changes in pupil diameter were recorded and measured online using custom routines kindly provided by Bahr, Kremkow, Sachdev and colleagues (Bergmann, 2019).

### *In vivo* juxtacellular recordings in POm in awake mice

To perform juxtacellular recordings from awake mice, mice first required implantation of a head bar. Under isoflurane-anesthesia (~1-2 % in O2, vol/vol), mice were placed in a stereotaxic frame (Narishige). Eye ointment was applied to the eye to prevent dehydration while body temperature was maintained at ~36 °C and the depth of anesthesia monitored throughout the experiment. Mice received a subcutaneous injection of lidocaine (1 %, wt/vol) and meloxicam (3 mg / Kg) was injected (i.p.) for analgesia and anti-inflammatory action. The skin at the surface of the skull was disinfected with ethanol 70% and betadine before to be surgically extracted and leaving the skull exposed. A head bar was then attached to the skull for head-fixation using dental cement (C&B metabond, Parkell Inc). Mice were allowed to recover in their home cage for 6 days before to the start of habituation to head fixation. During the habituation period, mice were handled and acclimatized to the behavioral setup, being head restrained for incremental periods of time. Once fully habituated (approximately after 1 week), mice were able to stay head-fixed for at least 90 min, during which they were also randomly presented with the tactile stimulus to the contralateral forepaw (500 ms, 200 Hz) to prevent startle response during recording. Once habituated, a craniotomy was performed, centered at coordinates: RC, 1.7 mm; ML, 1.68 mm; from bregma and the dura, under isoflurane-anesthesia (~1-2 % in O2, vol/vol). Meloxicam (3mg / Kg) was injected (i.p.) for additional post-operative analgesia and anti-inflammatory action. The craniotomy was then submerged with a mixture of normal rat ringer/agarose (2%) and was covered with silicone (Smooth-On, Inc.). Mice were placed back to their home cage for 2 hours for recovery prior to electrophysiological recordings. Following recovery, mice were headfixed, and the protective silicon removed to expose the craniotomy. A glass filament pipette (4-10 MΩ) filled with normal rat ringer was inserted into the brain at 10° angle to a depth of 2.57 – 3.00 mm depth at maximum pressure. Test pulses (−10 mV, 20 ms) were applied to the pipette in voltage clamp mode and the positive pressure was decreased to 25-35 mbar and the pipette advanced at steps of 1μm until a cell was encountered. Cell-attached juxtacellular recordings were performed using a differential amplifier (BVC-700A, Dagan). Custom written software in Igor Pro (WaveMetrics) was used for acquisition at 50 kHz in a voltage clamp mode. Mice behavior was monitored with an infrared sensitive camera (Microsoft lifecam) and recordings of trials where the forepaw was not in contact with the stimulator were discarded. The recording sites in each mouse were reconstructed posthoc to ensure targeting of the POm nucleus. The fluorescent protein tetramethyl rhodamine was included in the recording pipette and pressure ejected into the recording region after each experiment. The location of recordings were therefore identified by either directly visualizing fluorescent cells or visualization of the bolus load. Recordings of 1 out of the 5 mice were discarded as the pipette location were not found in POm.

### *In vivo* whole cell recordings and photo-stimulation of POm axons in forepaw S1

Approximately three weeks after stereotaxic injection of AAV1.hSyn.ChR2(H134R)-eYFP.WPRE.hGH (60 nl, see virus injection), mice were initially anaesthetized with isoflurane (~3 % in O2, vol/vol,) before urethane anesthesia (1.3 g/kg of body weight in 20 % saline, Sigma) was administered i.p. Body temperature was maintained at ~36 °C and the depth of anesthesia was monitored throughout the experiment and, when necessary, anesthesia was topped-up with 10 % of the initial urethane dose. Once anesthetized, lidocaine (1 %, wt/vol,) was injected around the surgical site and the mouse head was stabilized in a stereotaxic frame by a head-plate attached to the skull with dental cement (C&B metabond). A craniotomy was then performed above the forepaw area of the primary somatosensory cortex (~1.5 × 1.5 mm square, centered at coordinates: RC, 0 mm; ML, −2.3; from bregma) and the area was then submerged with rat ringer solution containing (135 mM NaCl, 5.4 mM KCl, 1.8 mM CaCl2, 1 mM MgCl2, 5 mM HEPES). In vivo whole cell patch recordings were obtained from L2/3 pyramidal neurons using pipettes with a tip resistance between 6 - 9 MΩ, filled with intracellular solution containing (in mM): 115 K gluconate, 20 KCl, 10 HEPES, 10 phosphocreatine, 4 ATP, 0.3 GTP adjusted to pH 7.3-7.4 with KOH. Recordings were performed from the soma using a differential amplifier (BVC-700A, Dagan) and were filtered at 10 kHz. Test pulses (10 mV,10 ms) were applied to the pipette in voltage clamp and using maximal pressure the pipette was inserted at 30° degree-angle into the forepaw area of S1 to a vertical depth of 200 μm after which the positive pressure was decreased to 25-35 mbar and the pipette advanced at steps of 1μm until a cell was encountered (initial access resistance following whole-cell were typically ~50 MΩ). Because the in vivo recordings were performed blind, pyramidal neurons were identified according to their voltage response to current steps. Custom written software in Igor Pro (WaveMetrics) was used for acquisition and no correction was made for the junction potential between the bath and pipette solutions. On occasion, holding current was applied to the neuron via the whole-cell recording pipette (~50-200 pA) to encourage the neuron to fire more action potentials required for analysis. Photo-stimulation of POm axonal terminals expressing ChR2 was achieved by coupling a LED light to a 5x objective (Olympus) to obtain a field of focus roughly the size of the craniotomy (1.5-2 mm in diameter). The objective was focused at the cortical surface and a LED light generator (OptoLED Light Source, CAIRN) was used to deliver 6 brief light pulses (10 Hz, 470 nm, 60 mW) at 10 ms duration which reliably activates neurons (Herman et al., 2014). Stimuli were designed and triggered by custom built software in Igor Pro (WaveMetrics). Forepaw stimulation (500 ms, 200 Hz) was provided by a stepper motor (Precision Microdrives) coupled to a rod onto which the animal forepaw rested. POm axons in forepaw S1 and the contralateral forepaw were stimulated in isolation or simultaneously. Interleaved stimuli were delivered at low frequency 0.1 Hz to prevent habituation or sensitization effects.

### Pharmacology

Pharmacological modulation of cortical GABA_A_ was performed in a cohort of mice by local application to the exposed surface of gabazine (10μM, Tocris). Typically, the effect of the drug was observable over the time course of 4–5min, and remained stable thereafter.

### *Ex vivo* whole cell recordings and photo-inhibition of POm neurons by ArchT activation

Mice (P40 – P45) previously injected with ArchT in the POm (>14 days prior) were anaesthetized with isoflurane (3 – 5 % in 0.75 L/min O_2_) before decapitation. The brain was then rapidly transferred and cut in an ice-cold, oxygenated solution containing (in mM): 110 choline chloride, 11.6 Na-ascorbate, 3.1 Na-pyruvate,, 26 NaHCO_3_, 2.5 KCl, 1.25 NaH_2_PO_4_, 0.5 CaCl_2_, 7 MgCl_2_ and 10 D-Glucose (sigma). Coronal slices of the POm (300 μm thick) were cut with a vibrating microslicer (Leica Vibratome 1000S) and incubated in an incubating solution containing (in mM): 125 NaCl, 3 KCl, 1.25 NaH2PO_4_, 25 NaHCO_3_, 1 CaCl_2_, 6 MgCl_2_ and 10 D-Glucose at 35 °C for 20 minutes, followed by incubation at room temperature for at least 30 minutes before recording. All solutions were continuously bubbled with 95%O_2_/5%CO_2_ (Carbogen). Whole-cell patch clamp somatic recordings were made from visually identified pyramidal neurons using DIC imaging. During recording, slices were constantly perfused at ~1.5 ml/min with carbogen-bubbled artificial cerebral spinal fluid (ACSF) containing (in mM): 125 NaCl, 25 NaHCO_3_, 3 KCl, 1.25 NaH_2_PO_4_, 1.2 CaCl_2_, 0.7 MgCl_2_ and 10 D-Glucose maintained at 30-34 °C. Patch pipettes were pulled from borosilicate glass and had open tip resistance of 5-7 MΩ filled with an intracellular solution containing (in mM):135 potassium gluconate, 70 KCl, 10 sodium phosphocreatine, 10 HEPES, 4 Mg-ATP, 0.3 Na_2_-GTP and 0.3% biocytin adjusted to pH 7.25 with KOH. Photoinhibition of POm neurons was achieved by shining a 565 nm LED light (1 s) onto the slice surface during somatic current injection steps (2s). Firing rates before and during light application were quantified and compared to the sae time period of the current step injection when no light was applied (Fig. S5).

### Histology

At completion of each experiment, mice were transcardially perfused with phosphate buffer (PB 0.1M) and 4% paraformaldehyde (PFA) solution. Brains were collected and post fixed overnight (~12 hrs.) in 4 % PFA at 4 °C before being cut into 200 μm coronal slices using a vibratome (Leica VT1000 Automated Vibratome) and mounted on glass slides using mounting medium containing nuclear staining dye DAPI (Fluoroshild, Sigma). Images of the brain slices were acquired using wide-field fluorescent microscopy (Zeiss Axio Imager 2). Images were taken such that excitation light (EYFP, 555 nm; DAPI, 430 nm) was optimized below the maximum pixel saturation value for each fluorophore. To evaluate virus (GCaMP6f, ArchT, ChR2, tetramethyl rhodamine) expression profiles in the POm complex, images of brain sections were registered to the corresponding coronal plates of the Paxinos mouse brain atlas (Paxinos and Franklin, 2001). Data from out of target injections or failed viral expression were removed from further analysis.

## Data analysis and statistical methods

### Ca^2+^ data

All analysis was performed using ImageJ and custom written routines in Matlab or Python 3. Horizontal and vertical drifts of imaging frames due to animal motion were corrected by registering each frame to a reference image based on whole-frame cross-correlation. The reference image was generated by averaging frames for a given field of view (FOV) in which motion drifts were minimal. Region of interests (ROIs) of axonal shafts or buttons were manually selected using the standard deviation of the entire imaging session (~ 6000 – 8000 frames). ROIs were selected so that each ROI represented a single POm axon. To calculate the baseline fluorescence (F_0_) for each ROI, first the average baseline florescence intensity (across 60 frames prior to stimulus onset, 2 seconds) of each trial was taken. Second, the rolling median of these average baseline values was measured and used as F_0_. Fluorescence traces are expressed as relative fluorescence changes, Δ*F/F* = *(F – F_0_)/F_0_*. Only Ca^2+^transients which were greater than 2x the baseline standard deviation *(F_0_ + (2x s.d.))* and above the threshold for a period longer than 200 ms were selected. The onset of a Ca^2+^ transient was defined as the time point at which a transient crossed the detection threshold *((F_0_ + (2x s.d.))*. Average Ca^2+^ transient probability was measured as 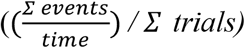. The peak amplitude (ΔF/F) was measured as the local maxima between the event onset and offset (i.e. when the falling edge of the transient crossed the threshold again). The duration (ms) of a Ca^2+^ transient was calculated as the time between the event onset and offset. Three behaviorally relevant epochs were selected for Ca^2+^ transient analysis (1 second duration each) for *spontaneous activity* (−2 to −1 sec, relative to stimulus onset); for *evoked activity* (0 to +1 sec, relative to stimulus onset) and for *reward activity* (0 to +1 sec, relative to reward delivery). For probability comparisons, all ROIs were used, while only the subset of ROIs (i.e. axons) with detectable events (greater than the threshold) were used to measure amplitude and duration. This determines the difference in number of axons used for each analysis. For direct comparison of spontaneous, response and or reward Ca^2+^ amplitude and duration, the subset of active axons with detectable Ca^2+^ events during both these behavioral epochs were used.

### Electrophysiology

During whole-cell recordings of L2/3 pyramidal neurons, custom-written Igor software (WaveMetrics) were used for both acquisition and analysis. A cell was selected for analysis if both ChR2-evoked subthreshold responses (i.e. EPSPs) and tactile-evoked subthreshold responses could be reliably measured. The average action potential frequency was measured for spontaneous firing and evoked firing (between −4 to −2 sec and 0 to +1sec, relative to stimulus onset, respectively). Custom-written Igor software (WaveMetrics) was used for both acquisition and analysis of juxtacellular recordings of POm neurons. Only stable neurons (that could last for 15 repetitions) and that present relatively stable action potentials according to their size were selected. Interspike intervals (ISI) were calculated during baseline and at the time of stimulus delivery (500 ms).

### Pupil tracking

Videos of pupil tracking and animal behavior were acquired and checked post hoc to remove potential artifacts due to sudden eyelid closing. Analysis of pupil dynamics were performed using a custom written algorithm in python. Briefly, pupil tracking for the entire session was split into single trials (11-sec duration) according to behavioral outcome. The average response profile was then calculated for each trial type for each mouse. Pupil dilation was monitored during a 4-sec baseline period preceding the beginning of each trial. The average peak diameter was measured as the local maxima of the average pupil response during the baseline epoch (−4 to 0 sec, relative to trial start), pre stim epoch (−3 to 0 sec, relative to stimulus onset) and post stim epoch (0 to +4 sec relative to stimulus onset).

### Behavior

The correct response rate was determined as d prime (the z transforms of HIT rate and FA rate d’ = z(H) - z(F)) or as the fraction of correct trials over the total number of trials (HIT trials + correct rejection trials)/(stimulus trials + catch trials). The behavioral effects of POm photo-inactivation were quantified by comparing correct responses of photo-inactivation (LED-ON trials) vs. control (LED-OFF) trials, typically 150 each per experimental session. LED-ON trials and LED-OFF trials were randomly interleaved. The latency to first lick was calculated as the time of first lick occurrence after stimulus onset.

### Statistical analysis

All statistics were performed using Prism software. The *α* significant level was set at 0.05. Normality of all value distributions was assessed by Shapiro-Wilk test (*α* = 0.05). Standard parametric tests were used only when data passed the normality test (p > 0.05). Non-parametric tests were used otherwise. Only two-sided tests were used. Specific statistical tests used and sample sizes are shown in figure captions or text.

## References

Alloway, K.D., Smith, J.B., Mowery, T.M., and Watson, G.D.R. (2017). Sensory Processing in the Dorsolateral Striatum: The Contribution of Thalamostriatal Pathways. Front Syst Neurosci 11, 53.

Audette, N.J., Bernhard, S.M., Ray, A., Stewart, L.T., and Barth, A.L. (2019). Rapid Plasticity of Higher-Order Thalamocortical Inputs during Sensory Learning. Neuron 103, 277–291 e274.

Audette, N.J., Urban-Ciecko, J., Matsushita, M., and Barth, A.L. (2017). POm Thalamocortical Input Drives Layer-Specific Microcircuits in Somatosensory Cortex. Cereb Cortex 28, 1312–1328.

Bergmann, R., Dominiak, S., Bahr, V., Kremkow, J., Mashaat, M.A., Oraby, H., Sehara, K., Larkum, M.E., Sachdev, R.N. (2019). Coordination of behavior in an Air Track maze: Sequential movement of whiskers, the maze and eyes. Paper presented at: Society of Neuroscience Abstracts.

Casagrande, V.A., Sary, G., Royal, D., and Ruiz, O. (2005). On the impact of attention and motor planning on the lateral geniculate nucleus. Prog Brain Res 149, 11–29.

Deschenes, M., Veinante, P., and Zhang, Z.W. (1998). The organization of corticothalamic projections: reciprocity versus parity. Brain Res Brain Res Rev 28, 286–308.

Gambino, F., Pages, S., Kehayas, V., Baptista, D., Tatti, R., Carleton, A., and Holtmaat, A. (2014). Sensory-evoked LTP driven by dendritic plateau potentials in vivo. Nature 515, 116–119.

Guo, Z.V., Inagaki, H.K., Daie, K., Druckmann, S., Gerfen, C.R., and Svoboda, K. (2017). Maintenance of persistent activity in a frontal thalamocortical loop. Nature 545, 181–186.

Herman, A.M., Huang, L., Murphey, D.K., Garcia, I., and Arenkiel, B.R. (2014). Cell type-specific and time-dependent light exposure contribute to silencing in neurons expressing Channelrhodopsin-2. Elife 3, e01481.

Hong, Y.K., Lacefield, C.O., Rodgers, C.C., and Bruno, R.M. (2018). Sensation, movement and learning in the absence of barrel cortex. Nature 561, 542–546.

Jahanshahi, M., Obeso, I., Rothwell, J.C., and Obeso, J.A. (2015). A fronto-striato-subthalamic-pallidal network for goal-directed and habitual inhibition. Nat Rev Neurosci 16, 719–732.

Jones, E.G. (2007). The Thalamus., Vol 2.

Kepecs, A., Uchida, N., Zariwala, H.A., and Mainen, Z.F. (2008). Neural correlates, computation and behavioural impact of decision confidence. Nature 455, 227–231.

Kobak, D., Brendel, W., Constantinidis, C., Feierstein, C.E., Kepecs, A., Mainen, Z.F., Qi, X.L., Romo, R., Uchida, N., and Machens, C.K. (2016). Demixed principal component analysis of neural population data. Elife 5.

Komura, Y., Tamura, R., Uwano, T., Nishijo, H., Kaga, K., and Ono, T. (2001). Retrospective and prospective coding for predicted reward in the sensory thalamus. Nature 412, 546–549.

Lacefield, C.O., Pnevmatikakis, E.A., Paninski, L., and Bruno, R.M. (2019). Reinforcement Learning Recruits Somata and Apical Dendrites across Layers of Primary Sensory Cortex. Cell Rep 26, 2000–2008 e2002.

Li, N., Chen, T.W., Guo, Z.V., Gerfen, C.R., and Svoboda, K. (2015). A motor cortex circuit for motor planning and movement. Nature 519, 51–56.

Mease, R.A., Metz, M., and Groh, A. (2016). Cortical Sensory Responses Are Enhanced by the Higher-Order Thalamus. Cell Rep 14, 208–215.

Meyer, H.S., Wimmer, V.C., Hemberger, M., Bruno, R.M., de Kock, C.P., Frick, A., Sakmann, B., and Helmstaedter, M. (2010). Cell type-specific thalamic innervation in a column of rat vibrissal cortex. Cereb Cortex 20, 2287–2303.

Micallef, A.H., Takahashi, N., Larkum, M.E., and Palmer, L.M. (2017). A Reward-Based Behavioral Platform to Measure Neural Activity during Head-Fixed Behavior. Front Cell Neurosci 11, 156.

Moore, J.D., Mercer Lindsay, N., Deschenes, M., and Kleinfeld, D. (2015). Vibrissa Self-Motion and Touch Are Reliably Encoded along the Same Somatosensory Pathway from Brainstem through Thalamus. PLoS Biol 13, e1002253.

Oh, S.W., Harris, J.A., Ng, L., Winslow, B., Cain, N., Mihalas, S., Wang, Q., Lau, C., Kuan, L., Henry, A.M., et al. (2014). A mesoscale connectome of the mouse brain. Nature 508, 207–214.

Paxinos, G., and Franklin, K.J. (2001). The Mouse Brain in Stereotaxic Coordinates.

Poort, J., Khan, A.G., Pachitariu, M., Nemri, A., Orsolic, I., Krupic, J., Bauza, M., Sahani, M., Keller, G.B., Mrsic-Flogel, T.D., et al. (2015). Learning Enhances Sensory and Multiple Non-sensory Representations in Primary Visual Cortex. Neuron 86, 1478–1490.

Poulet, J.F., and Petersen, C.C. (2008). Internal brain state regulates membrane potential synchrony in barrel cortex of behaving mice. Nature 454, 881–885.

Quiquempoix, M., Fayad, S.L., Boutourlinsky, K., Leresche, N., Lambert, R.C., and Bessaih, T. (2018). Layer 2/3 Pyramidal Neurons Control the Gain of Cortical Output. Cell Rep 24, 2799–2807 e2794.

Reimer, J., Froudarakis, E., Cadwell, C.R., Yatsenko, D., Denfield, G.H., and Tolias, A.S. (2014). Pupil fluctuations track fast switching of cortical states during quiet wakefulness. Neuron 84, 355–362.

Saalmann, Y.B., and Kastner, S. (2011). Cognitive and perceptual functions of the visual thalamus. Neuron 71, 209–223.

Schmitt, L.I., Wimmer, R.D., Nakajima, M., Happ, M., Mofakham, S., and Halassa, M.M. (2017). Thalamic amplification of cortical connectivity sustains attentional control. Nature 545, 219–223.

Sherman, S.M., and Guillery, R.W. (1996). Functional organization of thalamocortical relays. J Neurophysiol 76, 1367–1395.

Stringer, C., Pachitariu, M., Steinmetz, N., Reddy, C.B., Carandini, M., and Harris, K.D. (2019). Spontaneous behaviors drive multidimensional, brainwide activity. Science 364, 255.

Takahashi, N., Oertner, T.G., Hegemann, P., and Larkum, M.E. (2016). Active cortical dendrites modulate perception. Science 354, 1587–1590.

Trageser, J.C., and Keller, A. (2004). Reducing the uncertainty: gating of peripheral inputs by zona incerta. J Neurosci 24, 8911–8915.

Urbain, N., Salin, P.A., Libourel, P.A., Comte, J.C., Gentet, L.J., and Petersen, C.C.H. (2015). Whisking-Related Changes in Neuronal Firing and Membrane Potential Dynamics in the Somatosensory Thalamus of Awake Mice. Cell Rep 13, 647–656.

Wilke, M., Mueller, K.M., and Leopold, D.A. (2009). Neural activity in the visual thalamus reflects perceptual suppression. Proc Natl Acad Sci U S A 106, 9465–9470.

Williams, L.E., and Holtmaat, A. (2019). Higher-Order Thalamocortical Inputs Gate Synaptic Long-Term Potentiation via Disinhibition. Neuron 101, 91–102 e104.

Wimmer, R.D., Schmitt, L.I., Davidson, T.J., Nakajima, M., Deisseroth, K., and Halassa, M.M. (2015). Thalamic control of sensory selection in divided attention. Nature 526, 705–709.

Xu, N.L., Harnett, M.T., Williams, S.R., Huber, D., O’Connor, D.H., Svoboda, K., and Magee, J.C. (2012). Nonlinear dendritic integration of sensory and motor input during an active sensing task. Nature 492, 247–251.

Yamashita, T., Vavladeli, A., Pala, A., Galan, K., Crochet, S., Petersen, S.S.A., and Petersen, C.C.H. (2018). Diverse Long-Range Axonal Projections of Excitatory Layer 2/3 Neurons in Mouse Barrel Cortex. Front Neuroanat 12, 33.

Yamawaki, N., and Shepherd, G.M. (2015). Synaptic circuit organization of motor corticothalamic neurons. J Neurosci 35, 2293–2307.

Yu, J., Gutnisky, D.A., Hires, S.A., and Svoboda, K. (2016). Layer 4 fast-spiking interneurons filter thalamocortical signals during active somatosensation. Nat Neurosci 19, 1647–1657.

Yu, J., Hu, H., Agmon, A., and Svoboda, K. (2019). Recruitment of GABAergic Interneurons in the Barrel Cortex during Active Tactile Behavior. Neuron 104, 412–427 e414.

Zhang, W., and Bruno, R.M. (2019). High-order thalamic inputs to primary somatosensory cortex are stronger and longer lasting than cortical inputs. Elife 8.

Zhou, H., Schafer, R.J., and Desimone, R. (2016). Pulvinar-Cortex Interactions in Vision and Attention. Neuron 89, 209–220.

