## Supplementary Figures for "Higher order thalamus encodes correct goal-directed action"

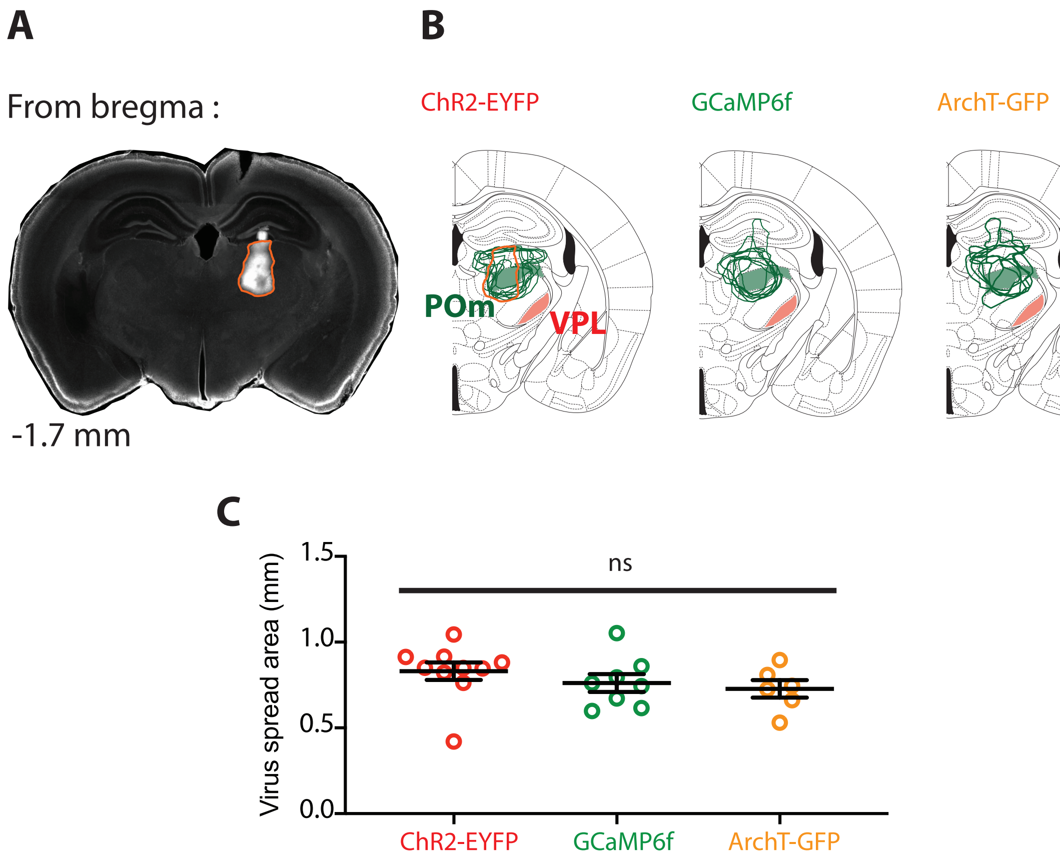


**Figure S1 | Targetting and spread of AAV injections in the POm nucleus.**
**(A)** Representative example of injection site in the mouse thalamus illustrating localized viral expression in the POm nucleus. **(B)** Overlay of the virus expression profile from the injection of ChR2-EYFP (left), GCaMP6f (middle) and ArchT-GFP (right) into the POm (green). The location- of the other thalamic nuclei which primarily projects to forepaw S1, the ventral posterolateral nucleus (VPL, red), is included for comparison. **(C)** Plot comparing the virus spread for the different injections.

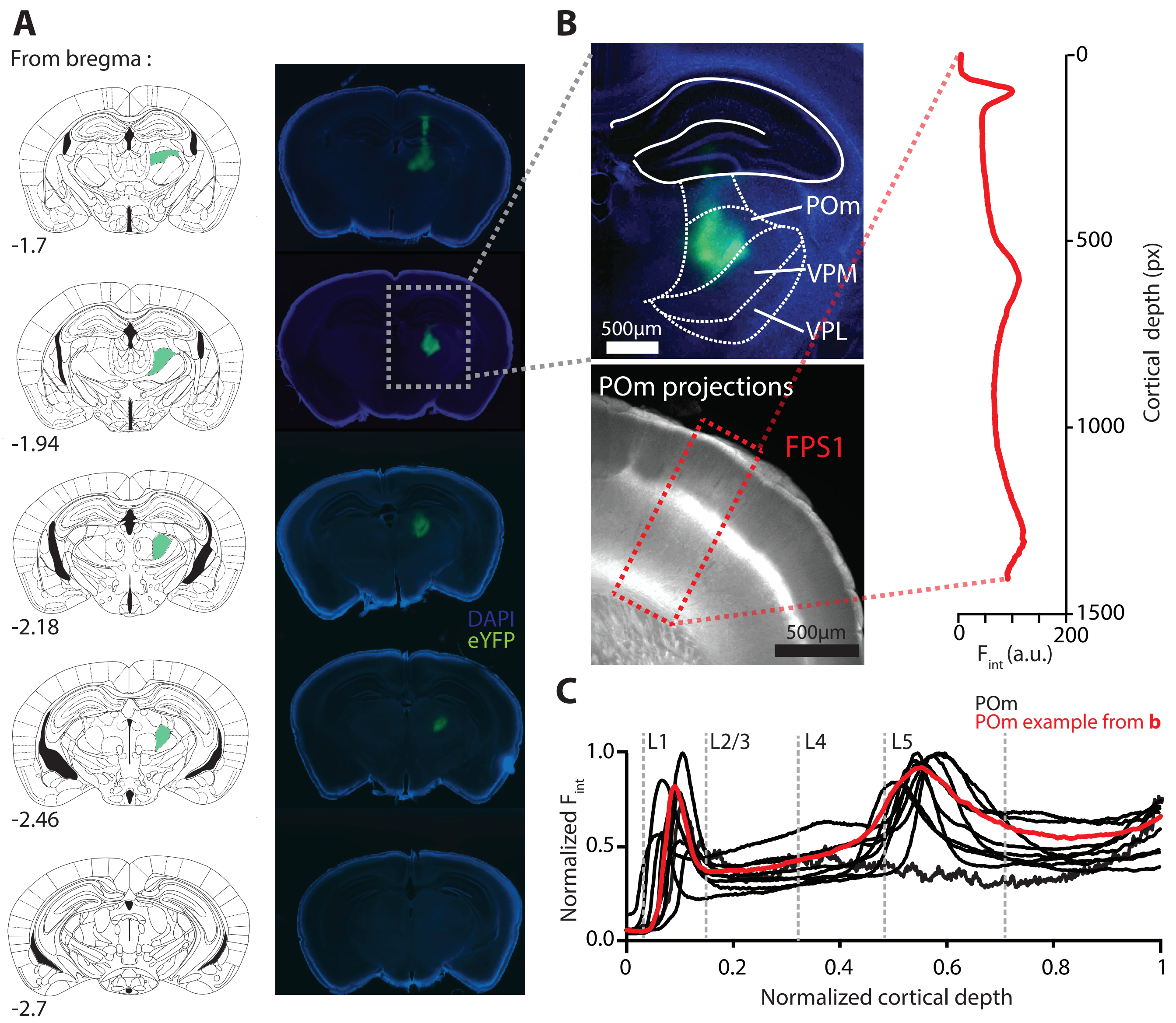


**Figure S2 | AAV mediated expression of ChR2-eYFP in the POm nucleus of the thalamus and its axonal projections in forepaw S1.**

**(A)** Representative example of ChR2-eYFP in the mouse thalamus illustrating localized viral expression in the POm nucleus. Left, outline of cortical slices at different rostra-caudal locations indicating the POm nucleus (green) from Paxinos and Franklin, 2001. Right, Corresponding brain sections showing the virus expression profile in the thalamus. **(B)** Top, enlarged fluorescence image showing ChR2-eYFP expression profile in POm nucleus from example in a. The location of the ventral posteromedial nucleus (VPM) and ventral posterolateral nucleus (VPL) are illustrated for comparison. Bottom, POm axonal projection profile in the forepaw area of S1 (FPS1) from example in a). Inset, Fluorescence intensity ($F_{int}$) measured as a function of cortical depth (in pixels (px)) in forpaw S1 (FPS1; rectangle). **(C)** Plot comparing the intensity profiles wihtin FPS1 of 10 different animals with injections localized in the POm nucleus.

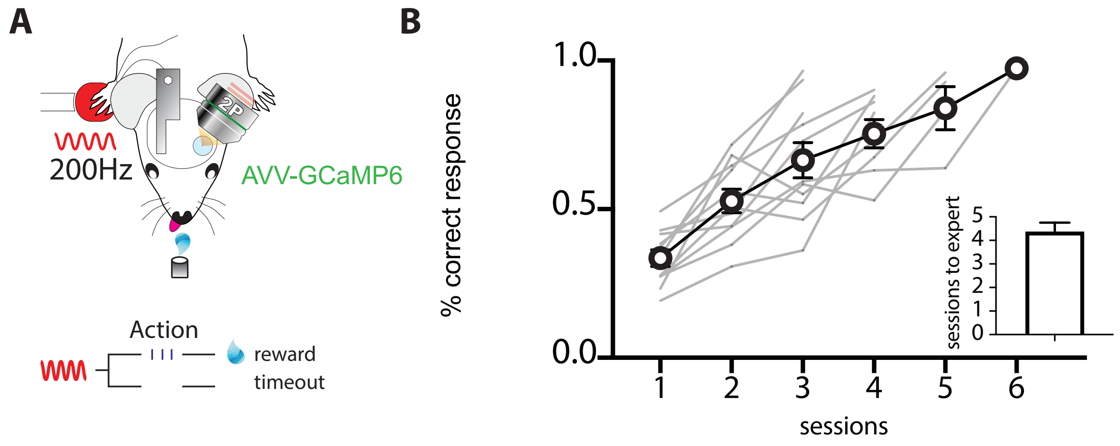


**Figure S3 | Mice rapidly learn the tactile goal-directed task.**
**(A)** Behavioral task design. Mice were habituated to head-restraint and trained to report the detection of a tactile stimulus (200 Hz, 500 ms) by licking a reward port. Correct responses were rewarded with sucrose water reward (10µl, 10% sucrose). Incorrect responses were punished with a timeout. **(B)** Performance of mice in the tactile goal-directed task improved in subsequent sessions, taking on average 4 days to reach expert performance (>80% correct; n = 11 mice)

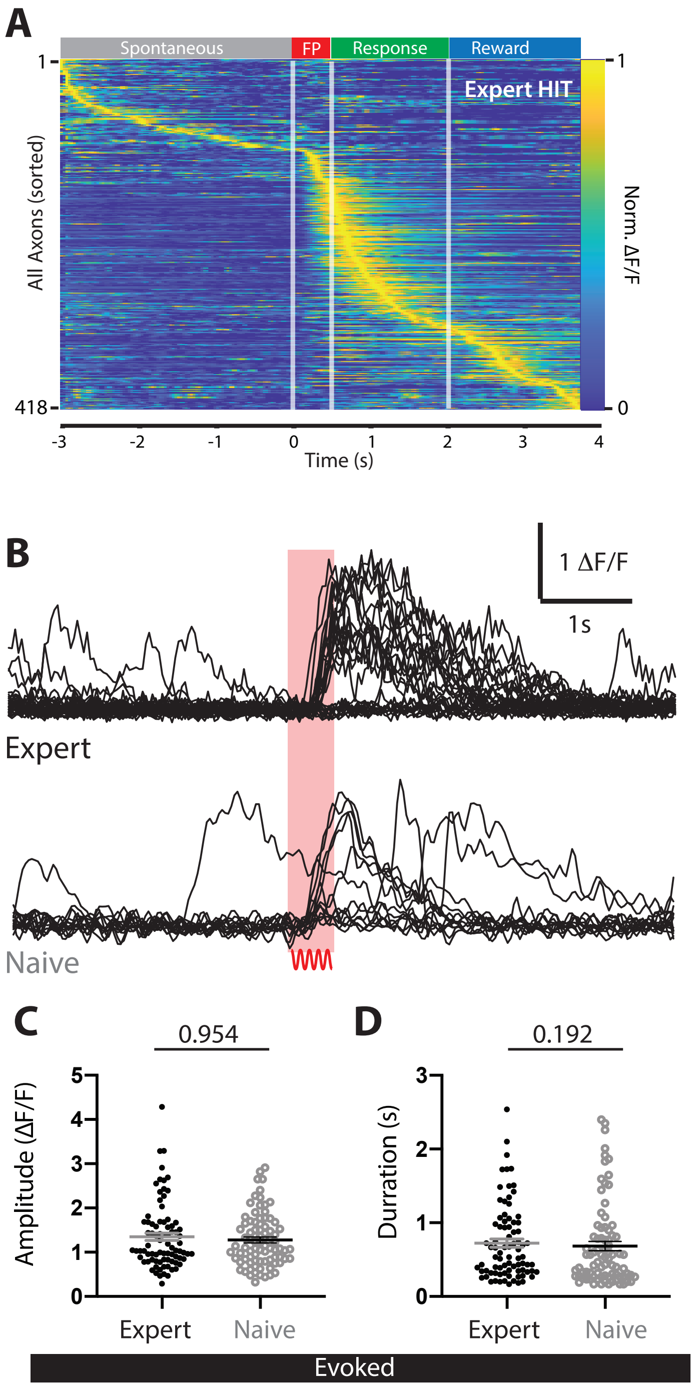


**Figure S4 | Tactile-evoked activity of POm axons projecting to forepaw S1 in naive and expert mice.**

**(A)** $Ca^{2+}$ activity pattern il all axons recorded (n = 418) during expert HIT responses. **(B)** $Ca^{2+}$ transients from POm axons projecting to forepaw S1 (top) during expert performance in a tactile goal-directed task and (bottom) in response to tactile stimulus to the contralateral forepaw in naive mice. Timing of stimulus indicated by red bar. **(C)** Peak amplitude of evoked $Ca^{2+}$ transients in expert (black) and naive (grey) mice. **(D)** Duration of evoked $Ca^{2+}$ transients in expert (black) and naive (grey) mice. Average amplitudes and durations for Expert were randomised and a sample of equal size to Naive was used for statisitical analysis. Mann-Whitney test was performed for significance testing.

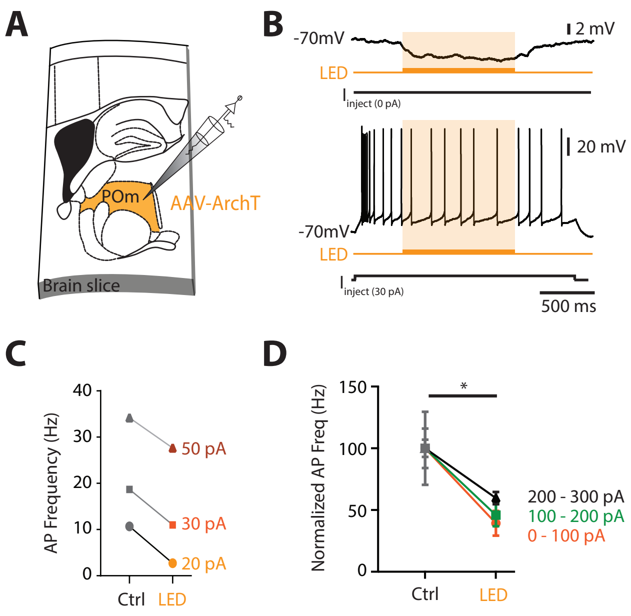


**Figure S5 | POm neurons are partially photo-inhibited by 590 nm LED.**
**(A)** The inhibitory opsin, archaerhodopsin (ArchT) was unilaterally injected into the POm and after 10-14 days, the brain was sectioned into 300 μm thick slices. Wholecell patch clamp voltage recordings were performed in POm neurons expressing ArchT. **(B)** Voltage recording during 590 nm light exposure from a POm neuron at rest (top) and during a current step injection (30 pA; 2 s; bottom). **(C)** Photo-inhibition caused a decrease in evoked action potentials during different current steps (20 pA, 30 pA, 60 pA) for example in b). **(D)** The POm was partially inhibited during LED exposure. 0 - 100 pA step pulse, p = 0.031. 100 - 200 pA step pulse, p = 0.016; 200 - 300 pA step pulse, p = 0.004. Wilcoxon matched-pairs signed rank test.

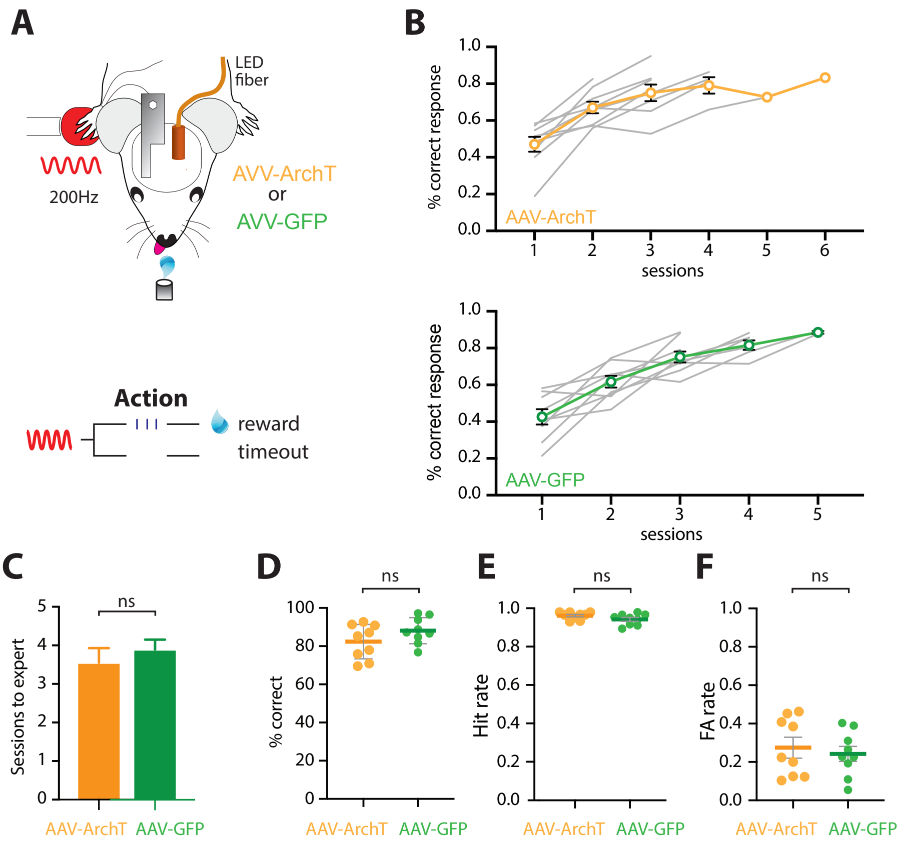


**Figure S6 | AAV injection did not alter goal-directed behavior.**

**(A)** Behavioral task design. Mice were habituated to head-restraint and trained to report the detection of a tactile stimulus (200 Hz, 500 ms) by licking a reward port. Correct responses were rewarded with sucrose water reward (10µl, 10% sucrose). Incorrect responses were punished with a timeout. **(B)** Performance in the tactile goal-directed task in mice previously injected with (top) the inhbitory opsin, AAV-ArchT (orange) and (bottom) the Green Fluorescent Protein control, AAV-GFP (green). Average with S.E.M, coloured line; individual mice, grey lines. **(C)** Number of training sessions required to reach expert performance (>80% correct). **(D)** Performance, measured as percent correct, in expert mice (p = 0.15; Unpaired t test). **(E)** Hit rate in expert mice (p = 0.11; Unpaired t test). **(F)** False alarm (FA) rate in expert mice (p = 0.21; Unpaired t test). Mice are considered expert when > 80% correct performance in the tactile goal-directed task. AAV-ArchT, orange, n = 9 mice; AAV-GFP, green, n = 9 mice. All data passed normality test (Shapiro-Wilk test).

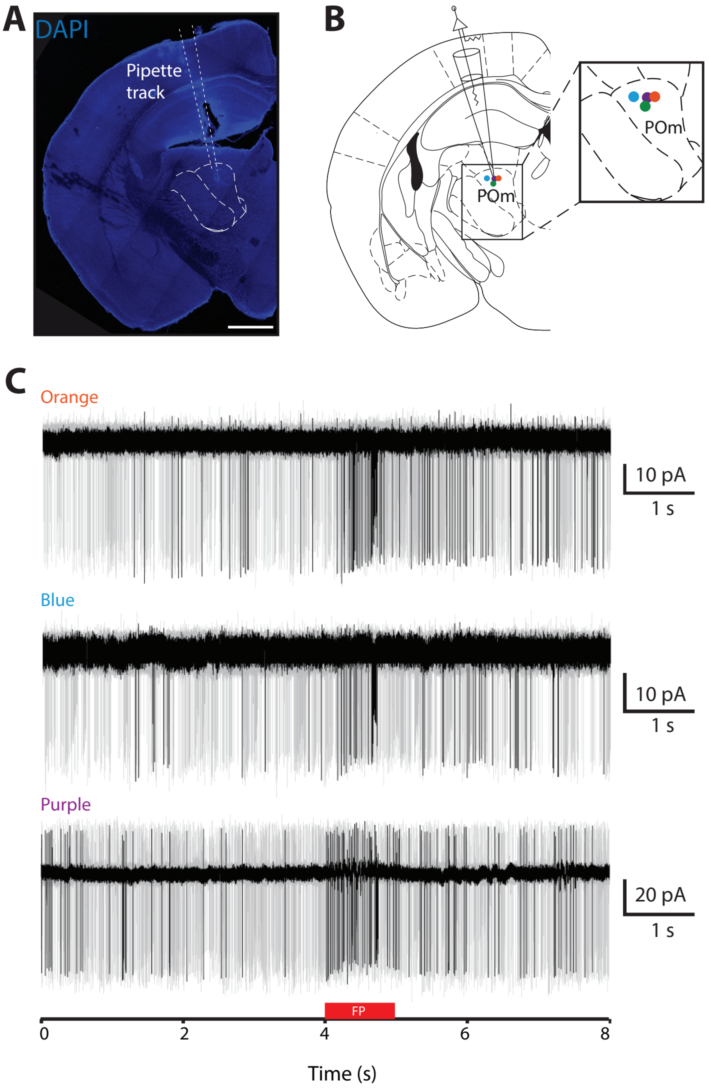


**Figure S7 | Juxtacellular cell attached recordings in POm neurons in awake mice.**
**(A)** DAPI stained brain slice illustrating the location of the recording pipette. **(B)** Post hoc reconstuction of the recording locations within the POm in 4 mice included in the analysis. **(C)** Typical examples of the response to tactile stimulus (500 ms; 200 Hz) delivered to the forepaw in POm neurons recorded at the orange (top), blue (middle) and purple (bottom) location within POm illustrated in b). Total POm neurons recorded at each location, orange, n = 4 neurons; blue, n = 2 neurons; purple, n = 4 neurons; green, n = 8 neurons. Time of stimulus, red bar.

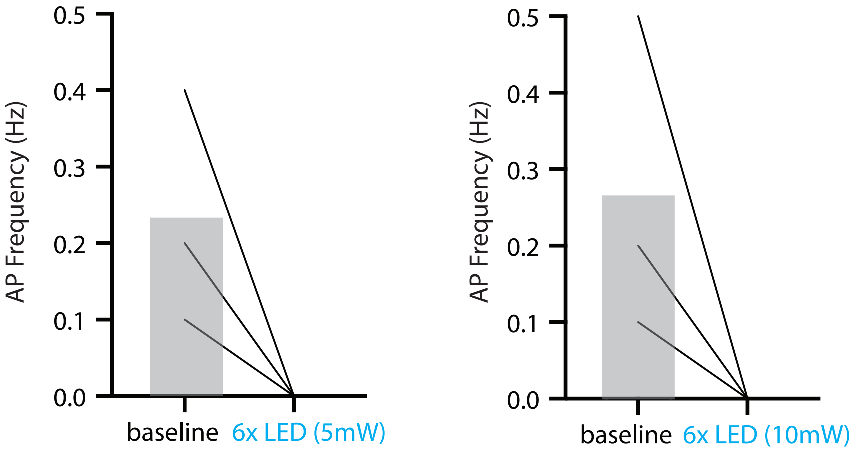


**Figure S8 | Activation of ChR2-expressing POm projections with different LED intensities inhibits action potentials in layer 2/3 pyramidal neurons within forepaw S1.**
Average action potential firing rate for layer 2/3 pyramidal neurons during baseline and sustained (6x, 10ms, 90 isi) POm photo-activation with (left) 5 mW and (right) 10 mW LED intensity.
